# Dynamic Transcriptomic Network Responses to Divergent Acute Exercise Challenges in Young Adults

**DOI:** 10.1101/2022.09.14.507939

**Authors:** Kaleen M Lavin, Zachary A Graham, Jeremy S McAdam, Samia M O’Bryan, Devin Drummer, Margaret B Bell, Christian J Kelley, Manoel E Lixandrão, Brandon Peoples, S. Craig Tuggle, Regina S Seay, Kendall Van Keuren-Jensen, Matthew J Huentelman, Patrick Pirrotte, Rebecca Reiman, Eric Alsop, Elizabeth Hutchins, Jerry Antone, Anna Bonfitto, Bessie Meechoovet, Joanna Palade, Joshua S Talboom, Amber Sullivan, Inmaculada Aban, Kalyani Peri, Timothy J Broderick, Marcas M Bamman

**Affiliations:** Healthspan, Resilience, and Performance, Florida Institute for Human and Machine Cognition, Pensacola, FL; UAB Center for Exercise Medicine, The University of Alabama at Birmingham, Birmingham, AL; Department of Cell, Developmental, and Integrative Biology, The University of Alabama at Birmingham, Birmingham, AL; Department of Biostatistics, The University of Alabama at Birmingham, Birmingham, AL; Birmingham VA Medical Center, Birmingham, AL; Cancer & Cell Biology, Translational Genomics Research Institute, Phoenix, AZ; Integrated Mass Spectrometry Shared Resource, City of Hope Comprehensive Cancer Center, Duarte, CA

## Abstract

Acute exercise elicits dynamic transcriptional changes that, when repeated, form the fundamental basis of adaptations in health, resilience, and performance. While moderate-intensity endurance training combined with conventional resistance training (traditional, TRAD) is often prescribed and recommended by public health guidance, high-intensity training combining maximal-effort intervals with intensive, limited-rest resistance training is a time-efficient alternative that may be used tactically (HITT) to seek whole body health benefits. Mechanisms of action of these distinct doses are incompletely characterized and have not been directly compared. We assessed transcriptome-wide responses in skeletal muscle and circulating extracellular vesicles (EVs) to a single exercise bout in young adults randomized to TRAD (n=21, 12M/9F, 22±3y) or HITT (n=19, 11M/8F, 22±2y). Next-generation sequencing captured small, long, and circular RNA in muscle and EVs. Analysis identified differentially expressed transcripts (|log_2_FC|>1, FDR≤0.05) immediately (h0, EVs only), h3, and h24 post-exercise within and between exercise doses. Additionally, all apparently responsive transcripts (FDR<0.2) underwent singular value decomposition to summarize data structures into latent variables (LVs) to deconvolve molecular expression circuits and inter-regulatory relationships. LVs were compared across time and exercise dose. TRAD generally elicited a stronger, more consistent transcriptional response than HITT, but considerable overlap and key differences existed. Findings reveal shared and unique molecular responses to divergent exercise stimuli and lay groundwork toward establishing relationships between protein-coding genes and lesser-understood transcripts that serve regulatory roles in response to exercise. Future work should advance the understanding of these circuits and whether they repeat in other populations or following other types of exercise/stress.

**NEW AND NOTEWORTHY:** We examined small and long transcriptomics in skeletal muscle and serum-derived extracellular vesicles before and after a single exposure to traditional combined exercise (TRAD) and high-intensity tactical training (HITT). Across 40 young adults, we found more consistent protein-coding gene responses to TRAD, whereas HITT elicited differential expression of microRNA enriched in brain regions. Follow-up analysis revealed relationships and temporal dynamics across transcript networks, highlighting potential avenues for research into mechanisms of exercise response and adaptation.

## INTRODUCTION

Exercise is a powerful and effective master regulator of metabolic health, physical performance, and overall wellness throughout the lifespan. As our team has recently reviewed (19), regular exposure to exercise improves cardiorespiratory fitness, neuromuscular function, cognitive function, metabolic efficiency, body composition, disease resilience, and healthspan, among a host of other benefits that continue to be identified. However, despite generally well-defined health and performance benefits, the optimal exercise “dose” continues to be an elusive research pursuit and likely varies based on age, fitness level, presence of comorbidities, other demographic confounders, and goal outcomes for training and/or overall health (22).

Exercise intensity and duration are the primary determinants of overall workload via their combined demands on neuromuscular activation patterns, cardiorespiratory demand, energy metabolism, and underlying molecular signaling required to support continued activity (19). Public health guidelines (e.g., the American College of Sports Medicine) generally encourage a combination of regular resistance and endurance training performed at a moderate-to-high intensity. In contrast to this more traditional (TRAD) approach, an increasingly popular modality both in the public (e.g., CrossFit, Tabata) and among military personnel (High Intensity Tactical Training, or HITT (13)), exists in the form of a circuit of movements performed at maximum velocity for brief time intervals with some movements performed explosively (i.e. maximum power) against bodyweight and/or added resistance. A range of functional and time-saving benefits are associated with HITT training, and some evidence indicates that the long-term physiological benefits are similar to TRAD training with regard to aerobic fitness (30), energy metabolism (28), and even longevity (50). However, whether these benefits arise through similar molecular mechanisms is not known. A better understanding of the so-called molecular circuits linked to TRAD vs. HITT responses may help guide exercise prescription for individuals, including identification of those whose intrinsic molecular phenotype suggests benefiting more from one dose than the other (22).

Mechanistic insight into the intensity-dependent molecular response to exercise is likely to provide valuable guidance towards optimizing and personalizing exercise dosing. The purest comparison may be best achieved in the exercise-naïve state following a single exposure to a relatively unaccustomed bout of exercise (15). Following acute exercise exposure, transient changes in gene expression form the basis of long-term adaptations at the protein level (33, 36), and therefore are most likely to lead to the myriad structural and functional changes in skeletal muscle and other organs and tissues. Accordingly, the current understanding of exercise transcriptomics is mostly limited to the subset of transcripts that encode protein (i.e., mRNA) and their ontological relationships to biological pathways via curated resources for annotation (16, 52). However, the vast majority of transcripts do not encode protein and are instead classified across a wide range of other “biotypes” that may serve a variety of signaling, regulatory, or otherwise poorly-understood cellular roles. For example, regulatory transcripts such as microRNAs (miRNAs) are a class of post-transcriptional modifiers that have been recognized to play important roles in exercise response and adaptation (10, 32). While produced in most tissues, miRNAs are commonly packaged into extracellular vesicles (EVs) and circulated to peripheral tissues (9), where they may affect a wide range of potential target genes. Additionally, a range of other non-coding biotypes of both small (e.g., piRNA, snRNA) and long (e.g., lncRNA, circRNA) nucleotide length is likely to exert regulatory effects on cellular processes including but not limited to transcription and translation (3, 12).

Bolstered by extensive research from our group (18, 20, 51) and many others (4, 37, 43), the “protein-coding gene” expression response to acute and chronic exercise provides a solid basis yet a still-incomplete picture of coordinated changes in the molecular environment. As such, this study examined the dose-dependent, coordinated transcriptomic responses to two exercise stimuli in a sex-balanced cohort in order to provide valuable insight across multiple knowledge gaps, including: (i) mechanistic clarity regarding acute molecular responses to HITT vs. TRAD, (ii) the relative coordinated temporal dynamics and potential regulatory response circuitry at the transcriptomic level, (iii) associations with established biological pathways linked to exercise adaptation. Profiling this response in healthy, young males and females not only strengthens the existing knowledge base but provides a foundation for more complex future studies investigating comparator populations (e.g., older adults, highly trained athletes, individuals with comorbidities, etc.), different exercise doses, and/or the effects of combined wellness strategies (e.g., exercise plus behavioral, nutritional, or pharmaceutical interventions).

## METHODS

### Participant Recruitment, Screening, and Randomization

For this investigation into the transcriptomic responses to acute exercise, 40 young adults (23M/17F, 22±3 y) were studied; representing a subset of participants enrolled in an exercise dose-response parent clinical trial (Department of Defense, Office of Naval Research grant: N000141613159, ClinicalTrials.gov Identifier: NCT03380923). Participants were recruited from the Greater Birmingham, Alabama area and surrounding communities. Inclusion/exclusion criteria and screening procedures are reported in NCT03380923. Briefly, all participants were aged 18-27y, generally healthy but not exercise-trained, had a body mass index (BMI) < 30, and were non-smokers. The trial was approved and monitored by The University of Alabama at Birmingham (UAB) Institutional Review Board for Human Use (IRB) and conducted in accordance with the Declaration of Helsinki. All participants provided written, informed consent to participate and allowed their biospecimens to be used for future research. A schematic of the general study design is presented in Fig. 1. Following consent and screening, participants were randomized to either traditional combined exercise training (TRAD, n=21, 12M/9F, 22±3 y) or high-intensity tactical training (HITT, n=19, 11M/8F, 22±2 y). Randomization was stratified by sex to ensure similar distributions of females and males in TRAD and HITT.

**Fig. 1:**
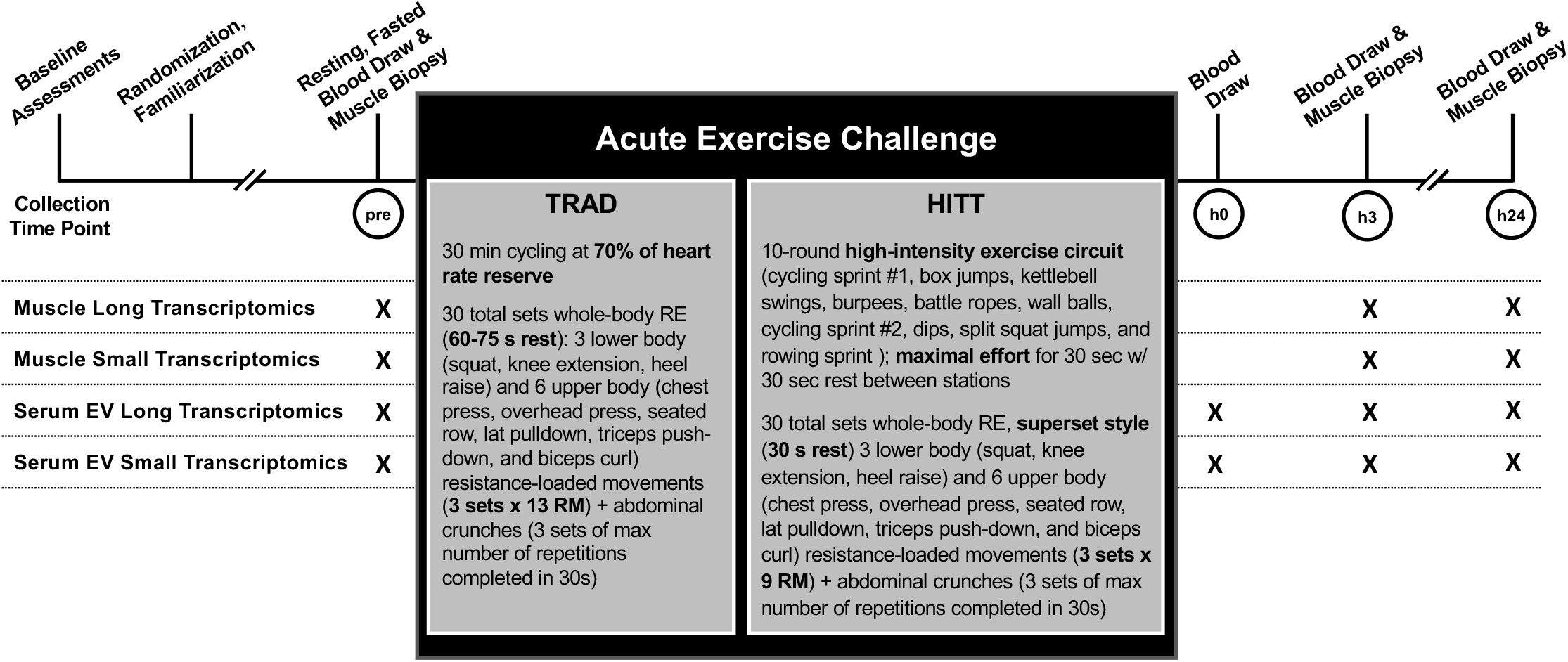
Schematic showing general study processes conducted around the acute exercise challenge and details for each bout. At the 978 indicated time points, participant underwent a muscle biopsy (pre-exercise, h3, and h24 post-exercise) and blood draw (pre-exercise,h0, h3, and h24 post-exercise) to be processed for extraction of serum extracellular vesicles (EV). Small and long transcriptomics were performed on RNA isolated from biospecimens collected at each time point.

### Phenotyping

Participants underwent several relevant phenotyping assessments to provide a detailed depiction of the cohort yielding the biospecimens and ultimately molecular response circuits. Baseline phenotyping included standardized and well-validated tests to evaluate body composition (dual-energy x-ray absorptiometry, DXA), cardiorespiratory fitness or aerobic power (VO_2_peak on a cycle ergometer), anaerobic power (30-second Wingate test), one-repetition maximum (1RM) upper and lower body dynamic strength on five movements, unilateral maximal voluntary isometric knee extension strength, MVC), and unilateral knee extension peak power. Methods for each of these tests is provided in detail elsewhere (2, 17, 40, 49).

### Familiarization

Participants completed four, progressive familiarization sessions prior to the acute exercise challenge. These familiarization sessions were designed to (i) ensure that all exercises were performed safely and correctly; (ii) dial in the target intensity and volume for the acute bout; and (iii) ensure that the acute bout indeed yielded a molecular exercise response in the untrained state as opposed to an overt tissue damage/inflammation/stress response in the truly naïve state. The four familiarizations followed a standardized progression in intensity and volume such that each participant’s first exposure to the full stimulus was the acute bout itself.

### Acute Exercise Challenge with Biospecimen Collection

All acute exercise bouts were performed between the hours of 0600 and 0900. Following an overnight fast (∼10h), individuals rested in a supine position for ∼30 minutes. A rested, fasting blood draw was obtained from an antecubital vein followed by a percutaneous muscle biopsy of the vastus lateralis using established methods previously described in detail (18, 30, 51). Muscle tissue was cleared of all visible connective tissue and fat, and ∼25-30 mg portions were immediately snap-frozen in LN_2_ for RNA isolation and other assays and stored at −80ºC until analysis. For histological analysis, a separate portion ∼3-4 mm in cross-sectional diameter was mounted on cork using OCT mixed with tragacanth gum and frozen in LN_2_ cooled isopentane. Blood was allowed to clot for 30 minutes and then spun at 2000 *x* g for 10 minutes at 4ºC to separate the serum fraction. Serum aliquots were stored at −80ºC until analysis.

Following the muscle biopsy, each participant performed the acute exercise challenge of the study arm to which they had been randomized (TRAD, HITT). **TRAD Prescription**. TRAD participants first performed 30 minutes of steady state cycle ergometry at an intensity targeting 70% of heart rate reserve (HRR), as calculated from resting HR and peak HR attained during the cycle ergometer-based maximal aerobic power (VO_2_max) test. This aerobic exercise was followed by 30 total sets of whole-body resistance exercise (RE) consisting of 3 lower body (squat, knee extension, heel raise) and 6 upper body (chest press, overhead press, seated row, lat pulldown, triceps push-down, and biceps curl) resistance loaded movements, along with abdominal crunches. For the nine loaded RE movements, 3 sets of each movement were completed with loads targeting an intensity of 13 repetition-maximum (13RM). Crunches were performed as 3 sets x maximum number of repetitions completed in 30 s per set. The total of 30 RE sets were performed with 60-75 s rest between sets. Given the focus on muscle biopsies of the vastus lateralis, the squat and knee extension were performed last in the sequence. **HITT Prescription**. In lieu of steady state aerobic exercise, HITT participants first performed a 10-round circuit of high-intensity, explosive exercises in the following order: cycling sprint #1, box jumps, kettlebell swings, burpees, battle ropes, wall balls, cycling sprint #2, dips, split squat jumps, and rowing sprint) each at maximal effort for 30 seconds with 30 seconds rest between stations. This was followed by whole-body RE with the same 10 movements x 3 sets each prescribed in TRAD. However, major differences in intensity and work-rest ratio were prescribed to HITT. Specifically, intensity of the loaded RE movements targeted 9RM resistance loads, supersets were performed consisting of 2 paired movements that activated antagonistic or independent muscle groups, and supersets were separated by only 30 s rest intervals. These differences were prescribed to contrast TRAD with a substantially more intense RE exposure in HITT. The prescriptions are summarized in Fig. 1. The entire protocol duration was ∼90 min for TRAD and ∼45 min for HITT(30).

No structured physical activity was allowed post-exercise through the h24 biospecimen collections. Immediately after the acute exercise bout, blood was collected (within ∼10 min, h0). Participants were then provided with a standardized protein shake and permitted to leave the exercise facility (with the activity restriction monitored by a wearable device) and returned for a post-exercise muscle biopsy and blood collection at h3. Subjects repeated an evening meal as similar as possible to the preceding night to reduce the influence of dietary variability within an individual, followed by an overnight fast (∼10h). The following morning, 24±2h from the cessation of the acute exercise bout, participants returned in the fasted state and underwent a final blood draw and muscle biopsy.

### RNA Isolation

RNA was isolated from both skeletal muscle and serum extracellular vesicles (EVs) for each study time point. Isolations were performed at UAB by two independent processors for each biospecimen type in batches balanced for sex and exercise group (TRAD/HITT), and all samples from a participant were always included in the same batch. For skeletal muscle, RNA was isolated from snap-frozen tissue (27.8±5.4 mg) using the Qiagen miRNeasy kit (Cat. No. 217004, Qiagen, Germantown, MD). The miRNeasy kit enables collection of all RNAs (including those <200 nt) and is therefore ideal for transcriptome-wide sequencing operations, capturing both long and small RNA. Muscle-derived RNA was confirmed to be of excellent quality via Agilent Bioanalyzer (Agilent, Santa Clara, CA) [average ± SD RNA integrity number (RIN) 8.5±0.4, and 28S/18S ratio 1.7±0.4].

EVs were isolated from approximately 1 mL serum. Any cryoprecipitates accumulated during freezing were cleared via centrifugation, and RNA was subsequently liberated from EVs using the exoRNeasy midi kit (Cat. No. 77144, Qiagen). The exoRNeasy kit is used in the only Clinical Laboratory Improvement Amendments (CLIA)-validated EV-based liquid biopsy tests and has been shown to have the highest reproducibility in replicates across multiple labs (45, 48).The kit has an affinity membrane that binds and isolates all EVs at once and eliminates contaminating RNA binding proteins, lipoproteins, and other RNA carriers by allowing them to pass though the membrane. Thus, RNA samples extracted from EVs were highly likely to be of sufficient quality for next-generation sequencing. Quant-iT Ribogreen RNA Assay (Cat. No. R11490, ThermoFisher, Waltham, MA) using the low-range concentration protocol confirmed good concentration across total EV-derived RNA samples (0.78 ± 0.41 ng/μL); however, due to the quantity of RNA necessary to determine RIN, bioanalysis was not performed. When total sample RNA exceeded 10 ng, 5 ng RNA was put into whole transcriptome library prep, and 5 ng RNA was put into small RNA library prep. For samples with less than 10 ng total RNA, half of the total RNA was put into whole transcriptome library prep and half was put into small RNA library prep.

### Next-Generation Sequencing

#### Preparation and Quality Control

RNA was prepared for sequencing and subsequently sequenced at Translational Genomics Institute (TGen, Phoenix, AZ). Due to differing optimal downstream sequencing approaches for detection and quantification of long and small transcriptomics, separate laboratory workflows were used to prepare fractions for long and small transcriptomic sequencing from the same original RNA aliquot. Total RNA from skeletal muscle was DNase-treated with TURBO DNA-free (Cat. No. AM1907, ThermoFisher), then purified with RNeasy MinElute Cleanup (Cat. No. 74204, Qiagen) using a modified MinElute Cleanup protocol to capture both small and long RNA species; namely, in step 2 of the standard MinElute protocol, 950 μL 100% EtOH was added to each sample instead of 250 μL. RNA was measured for quantity with Quant-iT Ribogreen RNA Assay (Cat. No. R11490, ThermoFisher) and quality with High Sensitivity RNA ScreenTape and buffer (Cat. Nos. 5067-5579 & 5067-5580, Agilent).

#### Long RNA Library Preparation and Next-Generation Sequencing

For each RNA sample, a uniquely dual-indexed, Illumina-compatible, double-stranded cDNA whole transcriptome library was synthesized from 10 ng (for skeletal muscle) or up to 5 ng (for serum EVs) total RNA with the SMARTer Stranded Total RNA-Seq kit v2 Pico Input Mammalian (Cat. No. 634418, Takara Bio, San Jose, CA) and SMARTer RNA Unique Dual Index Kit (Cat. No. 634452, Takara Bio). Briefly, this library preparation included RNA fragmentation, a 5-cycle Indexing PCR, ribosomal cDNA depletion, and enrichment PCR (12 cycles for skeletal muscle, 16 for serum EVs). RNA fragmentation incubation time and temperature were dictated by each sample’s RIN (when available) and %DV200 according to the library prep protocol. Each library was measured for size with Agilent High Sensitivity D1000 ScreenTape and buffer (Cat. Nos. 5067-5584 & 5067-5603, Agilent). Next, 1 μL of each library was combined into a non-equimolar pool, which was then measured for size via TapeStation and concentration via the KAPA SYBR FAST Universal qPCR Kit (Cat. No. KK4824, Roche, Wilmington, MA), diluted to 70 pM, then loaded into an iSeq flowcell cartridge (Cat. No. 20031371, Illumina, San Diego, CA) with a 1% v/v PhiX Control v3 spike-in (Cat. No. FC-110-3001, Illumina), and sequenced at 101 × 8 × 8 × 101 cycles. Passing filter cluster counts per library were generated from this data and used to make a re-balanced pool, which was subsequently measured for size and concentration, diluted to 2 nM with a 1% v/v PhiX Control v3 spike-in, denatured and further diluted, loaded into a NovaSeq 6000 flow cell cartridge (Cat. No. 20028313, Illumina), and sequenced at 101 × 9 × 9 × 101 cycles with standard workflow and a final flow cell concentration of 400 pM. Muscle cDNA libraries were sequenced to at least 40M read pairs (or 80M paired-end reads), and serum EV cDNA libraries were sequenced to at least 50M read pairs (or 100M paired-end reads).

#### Small RNA Library Preparation and Next-Generation Sequencing

Small RNA libraries were generated from 125 ng skeletal muscle RNA or 5 ng serum EV RNA using NEXTflex Small RNA Library Prep Kit v3 (Cat. No. NOVA-5132-23, Perkin Elmer, Waltham, MA). The manufacturer’s instructions were followed with the following modifications: (i) 3’ and 5’ adapters were diluted by half, (ii) adapter inactivation (protocol step C) was not performed, (iii) 2 μL adapter inactivation buffer was added after protocol step B, and (iv) an additional 0.5 μL of water was added to each well. 18 cycles of PCR amplification were used for serum EVs and 16 cycles for skeletal muscle. Libraries between 150 and 200 bp in size were excised and purified using 6% Tris/Borate/EDTA gels, followed by DNA cleaning, concentrating (Zymo, Irvine, CA), and eluting in 30 μL water. The quality and quantity of gel-purified libraries was assessed using Agilent 2100 Bioanalyzer with High-Sensitivity DNA chips. Equimolar amounts of libraries were pooled and quantified. Pooled libraries were then normalized and denatured at a working concentration range of 15 pM with 5% PhiX spike-in for flow cell cluster generation. Libraries were sequenced on the HiSeq 2500 with cluster generation on the cBot Flow Cell (Illumina). For sequencing small RNA, kits with specialized adaptors that have four randomized nucleotides on the end for ligation were used to reduce bias in small RNASeq (11, 38, 57).

#### Alignment

The Extracellular RNA Communications Consortium (exceRpt) pipeline was used for alignment of small RNA(42). This pipeline is specifically optimized and validated for extracellular RNAs with alignments in parallel for miRNA (miRBase, piRNABank, tRNA, etc.). Data were normalized to read depth. Long RNA was aligned using STAR v2.6+ (7) to GRCh38 and to additional lncRNAs from the highly curated LNCpedia (54) and GENCODE datasets. Circular RNA (circRNA) were identified from reads in the long transcriptomics data set using CIRI2 following mapping with Burrows-Wheeler Alignment (bwa), as described (14). Briefly, a 19-nucleotide overlap between the read and both sides of a backspliced junction was required to identify a circRNA. Single nucleotide polymorphism (SNP) calls available for the Arizona Study of Aging and Neurodegenerative Disease (AZSAND) Brain and Body Donation Program (BBDP) were leveraged to provide additional insight into genetics and exRNA detection. Count data were created using featureCounts (35). All FASTQ files and raw count matrices are available in the Gene Expression Omnibus (GEO) database under accession number GSE209880 (reviewers please use access token: *ibitiyeavbqjjir*).

### Computational Analysis

#### Data Processing

The general workflow, including the gene x sample matrix dimensions for each data stream throughout processing, is presented in Table 2. All downstream data processing and analysis was performed using R v4.1.2. Each transcriptomics data stream was analyzed independently and in parallel according to the same general computational pipeline, with minor adjustments made where necessary to optimize the workflow for small, long, and circular RNA in both skeletal muscle and serum EVs. Each data stream was initially cleaned in order to remove all corresponding post-exercise samples from individuals whose pre-exercise sample failed sequencing QC. The rationale for this approach was that a missing pre-exercise sample would preclude interpretation of an exercise-induced change at any other time point; however, if the outlier was a sample collected at any post-exercise time point, all corresponding samples were retained in order to maximize the statistical power of the data set.

**Table 1:** Demographic and Phenotyping Results

|  | TRAD | HITT |
| --- | --- | --- |
|  | n = 21 (12M, 9F) | n = 19 (11M, 8F) |
| Age (y) | 22 ± 3 | 22 ± 2 |
| Height (cm) | 174 ± 10 | 173 ± 6 |
| Weight (kg) | 74 ± 12 | 73 ± 11 |
| BMI (kg/m<sup>2</sup>) | 24 ± 3 | 24 ± 3 |
| *Body Composition* | n = 20 | n = 19 |
| Total Lean Mass (kg) | 50.5 ± 9.4 | 49.7 ± 9.9 |
| Total Fat Mass (kg) | 20.0 ± 8.0 | 20.0 ± 6.1 |
| Body Fat (%) | 27.0 ± 8.7 | 27.6 ± 7.6 |
| *One Repetition Maximum Strength (1RM)* | n = 20 | n = 18 |
| Overhead Press (kg) | 42 ± 14 | 40 ± 19 |
| Chest Press (kg) | 61 ± 25 | 61 ± 37 |
| Lat Pull Down (kg) | 46 ± 15 | 47 ± 18 |
| Squat (kg) | 85 ± 30 | 83 ± 37 |
| Knee Extension (kg) | 95 ± 32 | 88 ± 36 |
| *Unilateral Maximal Knee Extension* | n = 20 | n = 18 |
| Isometric MVC Peak Torque (Nm) | 228 ± 70 | 218 ± 80 |
| Isotonic Average Power (W) | 183.2 ± 81.3 | 210.5 ± 130.0 |
| Isotonic Peak Work (J) | 98.3 ± 26.7 | 100.9 ± 38.1 |
| Isotonic Total Work (J) | 449.7 ± 121.6 | 460.7 ± 169.7 |
| *Aerobic Power* | n = 21 | n = 19 |
| Absolute VO<sub>2</sub>max (L/min) | 2.6 ± 0.7 | 2.4 ± 0.6 |
| Relative VO<sub>2</sub>max (mL/kg/min) | 35.3 ± 7.1 | 32.7 ± 5.0 |
| RER at VO<sub>2</sub>max | 1.21 ± 0.06 | 1.24 ± 0.09 |
| *Anaerobic Power* | n = 19 | n = 18 |
| Absolute Peak Power (W) | 732 ± 235 | 680 ± 235 |
| Relative Peak Power (W/kg) | 9.68 ± 2.25 | 9.29 ± 2.02 |

TRAD: traditional combined exercise; HITT: high-intensity tactical training; MVC: maximum voluntary contraction; VO<sub>2</sub>max: maximal oxygen consumption; RER: respiratory exchange ratio.

**Table 2:** Computational Workflow and Dimensionality for Analyzed Data Streams

| Transcript-Level Differential Expression Analysis | Skeletal Muscle |  |  | Serum EVs |  |  |
| --- | --- | --- | --- | --- | --- | --- |
|  | long RNA | small RNA | circRNA | long RNA | small RNA | circRNA |
| Raw counts (transcripts x samples) | 94946 x 120 | 85542 x 120 | 39577 x 120 | 94946 x 148 | 85542 x 154 | 197605 x 148 |
| Filtering Steps: |  |  |  |  |  |  |
| 1. to remove participants missing pre-exercise sample | 94946 x 120 | 85542 x 120 | 39577 x 120 | 94946 x 140 | 85542 x 154 | 197605 x 140 |
| 2. by biotype (based on length at transcript maturity) | 86636 x 120 | 5398 x 120 | 39577 x 120 | 86636 x 140 | 5398 x 154 | 197605 x 140 |
| 3. for expression (cpm >0) in ≥25% of samples | 32295 x 120 | 2673 x 120 | 1812 x 120 | 62772 x 140 | 1313 x 154 | 1120 x 140 |
| Batch effect identification (adjusted with ComBat_seq if needed) |  | *adjusted* |  | - | - | - |
| Outlier (±6 MAD from median) identification | 32295 x 120 | 2673 x 120 | 1812 x 120 | 62772 x 133 | 1313 x 146 | 1120 x 140 |
| Filtering by expression (longRNA only) | 23329 x 121 | - | - | 57025 x 133 | - | - |
| *limma* differential expression analysis with voom correction |  |  |  |  |  |  |
| **Exploratory Transcriptomic Network Discovery** |  |  |  |  |  |  |
| FDR < 0.2 at any time point | 19444 | 1728 | 148 | 55325 | 772 | 310 |
| All biotype counts collapsed within tissue & filtered for shared samples |  | 21320 x 120 |  |  | 56407 x 128 |  |
| *PLIER* Singular Value Decomposition: |  |  |  |  |  |  |
| n LVs generated (n showing main effect of exercise) |  | 40 (23) |  |  | 32 (4) |  |

See text for complete workflow, including steps that do not affect data dimensionality. EVs: extracellular vesicles; cpm: counts per million; MAD: mean absolute deviation; FDR: false discovery rate; PLIER: Pathway-Level Information Extractor; LVs: latent variables

RNA biotypes were analyzed as part of the data stream corresponding with transcript length in mature form [e.g., we discarded fragments that mapped to protein-coding reads in the small data set, or incompletely processed pri-miRNA in the long RNA data set(56)]. Based on an available pre-defined threshold (41), biotypes with a mean nucleotide (nt) length >200 were treated as long RNA (e.g., protein-coding, lncRNA, antisense, and multiple subclasses of pseudogenes), and those <200 nt were considered small RNA (e.g., miRNA, rRNA, tRNA, snRNA, snoRNA, and vaultRNA). A total of 20 long RNA biotypes and 9 small RNA biotypes were retained following this filtering step.

Count data were filtered by expression based on non-zero expression in at least 25% of samples. Next, principal components analysis (PCA) was used to identify the influence of any technical confounders, such as processing and/or sequencing batch. Where identified, batch effects were adjusted using ComBat_seq via *sva* (58). After applying a log_2_cpm normalization corrected for within-subject variance, statistical outliers were identified using PCA. Samples with scores more than six median absolute deviations from the median for PCs 1 and 2 (together, contributing to ∼35-40% of between-sample variance) were flagged as statistical outliers. An approach similar to that described above was used to clean the data for data points detected as statistical outliers and related samples. As a result, there was imperfect overlap across the six data streams, which we allowed in order to maximize the statistical power of each independent analysis.

#### Transcript-Level Differential Expression Analysis

While all biotypes were retained in the transcript-level differential expression analysis, interpretation and visualization were limited to protein-coding transcripts and microRNAs, based on resources currently available for annotation of biological roles. The *limma* package (23) was used to perform linear mixed models to compare transcript expression in each group over time, as well as any potential interaction of group and time, as described (25). In order to calculate predicted variance-based weights for each feature, voom normalization was performed across all transcripts (24), and all steps within the pipeline were blocked using participant ID as a random effect in order to control for between-subject variability. Resultant differentially expressed genes of interest were identified based on dual criteria for fold-change [absolute log_2_fold-change(FC) >1, corresponding to a doubling of halving of expression] and Benjamini-Hochberg adjusted false discovery rate (FDR<0.05). These thresholds were uniformly applied to all gene sets to isolate salient and consistent patterns in differential transcript expression.

Differences between groups were qualitatively visualized for all protein-coding transcripts and miRNAs surpassing fold-change and significance thresholds defined above. Protein-coding transcripts (from long RNA) and miRNA (from small RNA) were grouped into contrasts based on directional changes per time point per group. Differentially expressed circRNA were not pursued due to the low number of differentially expressed transcripts and insufficient availability of data for biological annotation, leaving a total of four data streams (protein-coding and miRNA for both muscle and serum EV). Within each, “intersections” (i.e., overlaps between transcript sets) were identified for all contrasts using *upsetR* (5), then ranked according to the number of genes in each intersection. For the eight largest intersections, downstream gene annotation was performed on protein-coding genes to identify biological pathways using overrepresentation analysis in both Gene Ontology and Kyoto Encyclopedia of Genes and Genomes (KEGG) databases. Pathway annotations were filtered for unadjusted p<0.001.

#### Exploratory Transcriptomic Network Discovery

In order to identify potentially important regulatory relationships across all biotypes, any transcript with an FDR<0.2 vs. pre-exercise at any post-exercise time point was considered responsive to acute exercise and included in the discovery analysis (an example for h3 vs. pre-exercise in skeletal muscle and serum EVs of the TRAD group is shown in Supplementary Fig. 1). This threshold was applied in order to reduce noise from unresponsive transcripts while allowing for inclusion of transcripts strongly trending towards significance, so as not to limit the discovery potential of the analysis. Separately but in parallel, all responsive transcripts within muscle (n=21,320) and serum EVs (n=56,407) were pooled and prepared for downstream analysis using Pathway-Level Information Extractor (*PLIER*)(29). Tissues were handled separately due to the differential impact of exercise on body compartments [e.g., only ∼33% (18,586 of 56,407) of genes meeting criteria for exercise responsiveness in serum EVs met the criteria in muscle], as well as the incompatible number of subjects between the two data streams as a result of strict upstream filtering (e.g., 96 of 120 samples in muscle were represented in serum EVs).

The PLIER algorithm is built around a singular value decomposition (SVD) anchored to biological pathway annotations for genes, as previously described (29) and used by our research team (18, 20). Presently, however, prior knowledge information was not provided in order to allow all unannotated transcripts (and therefore all biotypes) to remain in the algorithm; thus, PLIER essentially performed unsupervised SVD clustering across all transcripts to summarize patterns in the data independently of transcript biotype and/or known biological roles. The resulting data structures are known as latent variables, or LVs (29) and represent the variation across all samples. Each transcript is given a *value* within the dimension of the newly-defined LV (matrix Z (29)), whereas each sample is given a *score* representing the LV (matrix B (29)). Based on the structure of each data stream, PLIER identified 40 LVs from muscle and 32 LVs from serum EV responsive transcripts.

Using the generated scores, LVs were tested across samples using a group x time analysis of variance (ANOVA) design approximately analogous to that applied during the transcript-level analysis, and a correction factor for the number of LVs was applied to all ANOVA results to identify LVs of interest (p<0.05/number of LVs). Where an LV reached this strict threshold (i.e., p<0.00125 in skeletal muscle and p<0.00156 in serum EVs), it was followed up with a post-hoc test to identify the contrast for which a main or interaction effect was detected. All post-hoc tests were adjusted internally using a Benjamini-Hochberg adjustment for multiple comparisons; no additional adjustment was performed for the total number of LVs. All LVs surviving post-hoc testing were pursued for downstream interpretation. In order to strategically reduce the number of features pertinent to each LV, top transcripts were defined as any with a loading value greater than half the loading value for the top transcript in the LV (e.g., if the top-loaded transcript had a value of 0.70, all transcripts with values <0.35 were excluded). Transcripts were grouped by biotype and visualized to represent the biological makeup of the LV, and the protein-coding fraction of the LV was annotated using an approach identical to that described above (Gene Ontology and KEGG overrepresentation analysis). When possible, pathway annotations were filtered for p<0.001; where this approach yielded nothing, nominally significantly related pathways (p<0.05) were presented. Finally, LVs were classified based on general temporal pattern independently of direction into: “early response” (i.e., different at h0 and/or h3 post-exercise), “late response” (i.e., different at h24 post-exercise), or “sustained response” (i.e., different at h0 and/or h3 point and not returned to pre-exercise by h24).

## RESULTS

### Transcript-Level Differential Expression Analysis

#### Skeletal Muscle Long RNA

Differential skeletal muscle long transcript expression patterns following the acute exercise challenge are shown in Table 3. Based on the imposed thresholds, a total of 811 long transcripts (54% protein-coding) were upregulated at h3 post-exercise in the TRAD group, whereas 425 long transcripts (also 54% protein-coding) were upregulated in HITT; of these, 352 (206 protein-coding) were shared. Conversely, 2316 long transcripts (24% protein-coding) were downregulated in TRAD and 726 (22% protein-coding) in HITT at h3 post-exercise, with 584 (108 protein-coding) in common. At h24 post-exercise, a total of 932 (77% protein-coding) transcripts were upregulated in TRAD, whereas 783 (73% protein-coding) transcripts were upregulated in HITT; of these, 481 (396 protein-coding) were shared. TRAD also showed downregulation of 1058 long transcripts (31% protein-coding), while 479 (37% protein-coding) were downregulated in HITT, and 341 (131 protein-coding) transcripts were shared. Figure 2 shows the patterns in protein-coding genes at h3 (a) and h24 (b) post-exercise with respect to the FDR and log_2_fold-change vs. pre-exercise in TRAD only, given the overall greater magnitude of the acute skeletal muscle long transcript expression response in TRAD. Green circles represent transcripts where a significant change in expression was also detected in HITT.

**Table 3:** Number of Differentially Expressed Transcripts in Skeletal Muscle

|  |  |  | **all long**<br>**RNA** | **protein-**<br>**coding** | **all small**<br>**RNA** | **micro**<br>**RNA** | **circular**<br>**RNA** |
| --- | --- | --- | --- | --- | --- | --- | --- |
| h3 vs. pre-exercise | up | TRAD | 811 | 434 | 52 | 37 | 4 |
|  |  | HITT | 425 | 230 | 2 | 2 | 0 |
|  | down | TRAD | 2316 | 546 | 16 | 15 | 7 |
|  |  | HITT | 726 | 160 | 0 | 0 | 0 |
| h24 vs. pre-exercise | up | TRAD | 932 | 721 | 100 | 41 | 1 |
|  |  | HITT | 783 | 575 | 77 | 21 | 0 |
|  | down | TRAD | 1058 | 324 | 8 | 5 | 2 |
|  |  | HITT | 479 | 176 | 3 | 3 | 0 |

All transcripts satisfied both $|\log_2FC| > 1$ and $FDR < 0.05$ .

**Fig. 2:**
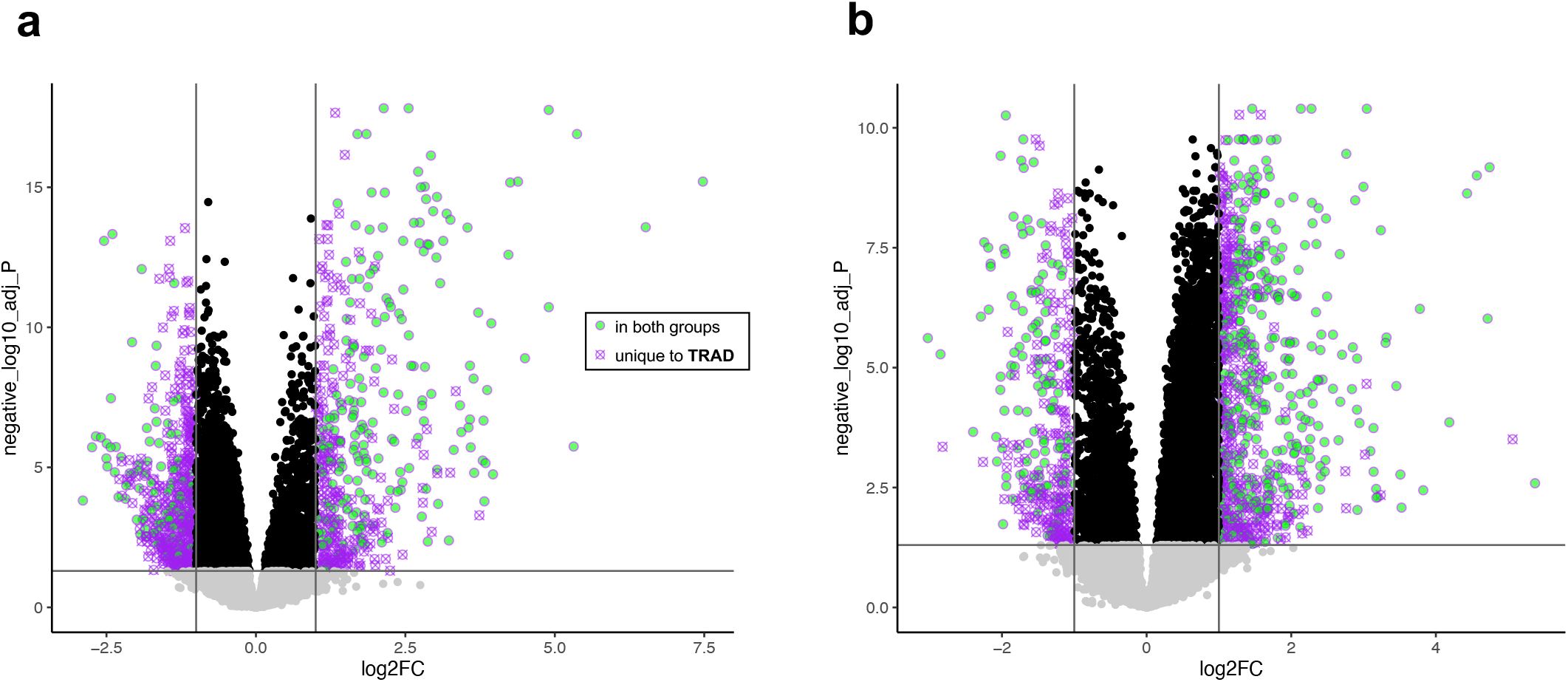
Volcano plot showing changes in protein-coding transcript expression **(a)** h3 and **(b)** h24 post-exercise in skeletal muscle of individuals performing TRAD exercise (|log_2_FC| >1, FDR<0.05). Transcripts meeting identical thresholds in HITT are represented as green circles, whereas those shown as crossed-out purple circles were only significant in TRAD (exact values plotted for log_2_FC and FDR correspond only to TRAD).

In order to establish insight into coordinated patterns in gene expression across time for both exercise doses, upsetR plots were constructed from the lists of up- and downregulated protein-coding transcripts at each time point vs. pre-exercise (Fig. 3a). The largest intersection contained 356 genes uniquely downregulated in TRAD at h3. Follow-up annotation revealed associations to Gene Ontology (GO) pathways including cell adhesion (GO:0007155, p=1.5×10^−7^), inflammatory response (GO:0006954, p=5.2×10^−7^), and cell activation (GO:0001775, p=8.5×10^−7^), among others. A common set of 282 genes upregulated in both groups at h24 post-exercise annotated primarily to metabolic processes related to glycerol-3-phosphate (GO:0006072; p=1.1×10^−5^), alditol phosphate (GO:0052646, p=2.3e^-5^), and carbohydrate (GO:0005975, p=7.3×10^−5^), among other pathways. These and further annotations for the top eight intersections (Fig. 3a), where available, are shown in Supplementary Table 1.

**Fig. 3:**
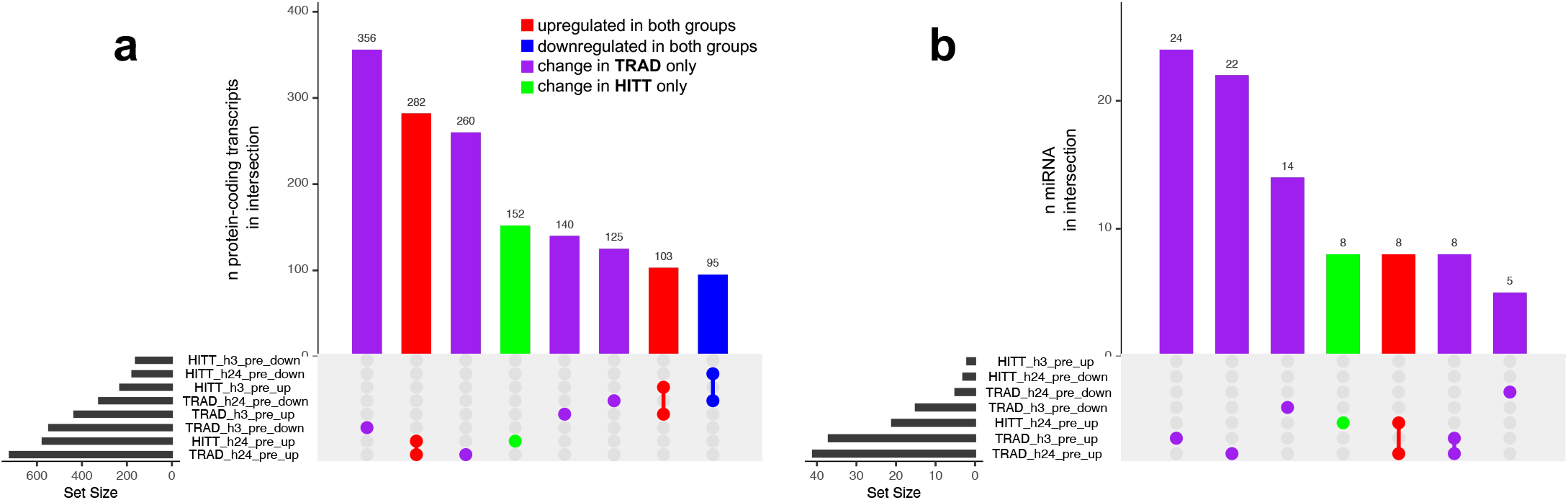
UpsetR plots indicating shared overlap (intersection) of directional transcript expression changes vs. pre-exercise in **(a)** protein-coding transcripts and **(b)** miRNA in skeletal muscle. Bar height represents the number of genes in each intersection, whereas dots represent comparison[s] in which transcripts were significantly up- or down-regulated vs. pre-exercise (|log_2_FC| >1 and FDR<0.05). Gene intersections are shown in red where upregulation was detected at any time point independently of group, in blue where downregulation was detected independently of group, in purple where a change was detected in TRAD only, and in green where a change was detected in HITT only. Gene ontology analysis for each intersection in panel **a** is available in Supplementary Table 1.

#### Skeletal Muscle Small RNA

At h3 post-exercise in TRAD, 52 small transcripts (71% miRNA) were upregulated, whereas 2 of these (both miRNAs) were upregulated in HITT. Conversely, 16 transcripts (94% miRNA) were downregulated in TRAD at h3, although none were significantly downregulated in HITT. At h24 post-exercise, a total of 100 small transcripts (41% miRNA) were upregulated in TRAD, whereas 77 small transcripts (27% miRNA) were upregulated in HITT; of these, 40 (11 miRNA) were shared. TRAD also showed downregulated expression of 8 small transcripts (5 miRNA), while 3 (all miRNA) were downregulated in HITT, though there was no overlap between groups. Supplementary Fig. 2 shows the patterns in miRNA at h3 (a) and h24 (b) post-exercise with respect to the FDR and log_2_fold-change vs. pre-exercise in TRAD only, with filled shapes representing transcripts where a significant change in expression was also detected in HITT. As shown in Fig. 3b, the small transcript response in TRAD was largely unique to this cohort, as the largest three intersections contained miRNA with unique responses in TRAD.

#### Skeletal Muscle Circular RNA

At h3 post-exercise in TRAD, 4 circRNAs were upregulated and 7 were downregulated. At h24 in TRAD, 1 circRNA was upregulated and 2 were downregulated. No circRNAs were significantly different from pre-exercise at any point in HITT. Given that the response in skeletal muscle circular RNA was overall more muted compared to the other data streams, these acute changes are not visually presented.

#### Serum EV Long RNA

Differential serum EV long transcript expression patterns following the acute exercise challenge are shown in Table 4. A total of 33,029 long transcripts (13% protein-coding) were upregulated at h0 post-exercise in TRAD, whereas 15,664 (perhaps notably, also 13% protein-coding) were upregulated in HITT. Of these, 14,141 (1,682 protein-coding) were common to both groups. Conversely, 259 long transcripts (81% protein-coding) were downregulated in TRAD and 223 (82% protein-coding) in HITT at h0 post-exercise, with 59 (55 protein-coding) in common. At h3 post-exercise, a total of 29,766 long transcripts (13% protein-coding) were upregulated in TRAD, whereas 20,629 (14% protein-coding) transcripts were upregulated in HITT; of these, 17,762 (2,370 protein-coding) were shared. TRAD also exhibited downregulation of 198 long transcripts (75% protein-coding), while 374 (83% protein-coding) were downregulated in HITT, and 156 (128 protein-coding) transcripts were shared. Finally, at h24 post-exercise, a total of 26,895 (12% protein-coding) transcripts were upregulated in TRAD, whereas 16,018 (14% protein-coding) transcripts were upregulated in HITT; of these, 13,310 (1716 protein-coding) were shared. TRAD also led to downregulation of 78 long transcripts (71% protein-coding), while 177 (78% protein-coding) were downregulated in HITT, and 59 (47 protein-coding) transcripts were common to both groups. Supplementary Fig. 3 shows the patterns in protein-coding transcripts at h0 (a), h3 (b), and h24 (c) post-exercise with respect to the FDR and log_2_FC vs. pre-exercise in TRAD only, with filled shapes representing where a significant change in expression was also detected in HITT.

**Table 4:** Number of Differentially Expressed Transcripts in Serum Extracellular Vesicles

|  |  |  | **all long**<br>**RNA** | **protein-**<br>**coding** | **all small**<br>**RNA** | **micro**<br>**RNA** | **circular**<br>**RNA** |
| --- | --- | --- | --- | --- | --- | --- | --- |
| h0 vs. pre-exercise | up | TRAD | 33029 | 4197 | 2 | 2 | 1 |
|  |  | HITT | 15664 | 2056 | 24 | 9 | 30 |
|  | down | TRAD | 259 | 211 | 0 | 0 | 0 |
|  |  | HITT | 223 | 183 | 0 | 0 | 6 |
| h3 vs. pre-exercise | up | TRAD | 29766 | 3967 | 4 | 0 | 0 |
|  |  | HITT | 20629 | 2975 | 8 | 2 | 2 |
|  | down | TRAD | 198 | 149 | 52 | 47 | 0 |
|  |  | HITT | 374 | 309 | 148 | 139 | 3 |
| h24 vs. pre-exercise | up | TRAD | 26895 | 3272 | 7 | 0 | 0 |
|  |  | HITT | 16018 | 2285 | 0 | 0 | 3 |
|  | down | TRAD | 78 | 55 | 50 | 49 | 0 |
|  |  | HITT | 177 | 138 | 126 | 121 | 1 |

All transcripts satisfied both $|\log_2FC| > 1$ and $FDR < 0.05$ .

As shown in Fig. 4a, the largest intersection contained 1,199 genes commonly upregulated at all post-exercise time points in both groups. GO analysis revealed these genes were primarily associated with protein-containing complex organization (GO:0043933, p=9.3×10^−10^), organelle organization (GO:0006996, p=7.8×10^−9^), and cellular component assembly (GO:0022607, p=1.1×10^−8^), among others. The next largest intersection contained 839 genes upregulated at every post-exercise time point in TRAD only. These genes primarily annotated to biological processes such as plasma membrane-bounded cell projection organization (GO:0120036, p=3.6×10^−6^) and neuron development (GO: 0048666, p=1.1×10^−5^). Other annotations, where available, are presented in Supplementary Table 2.

**Fig. 4:**
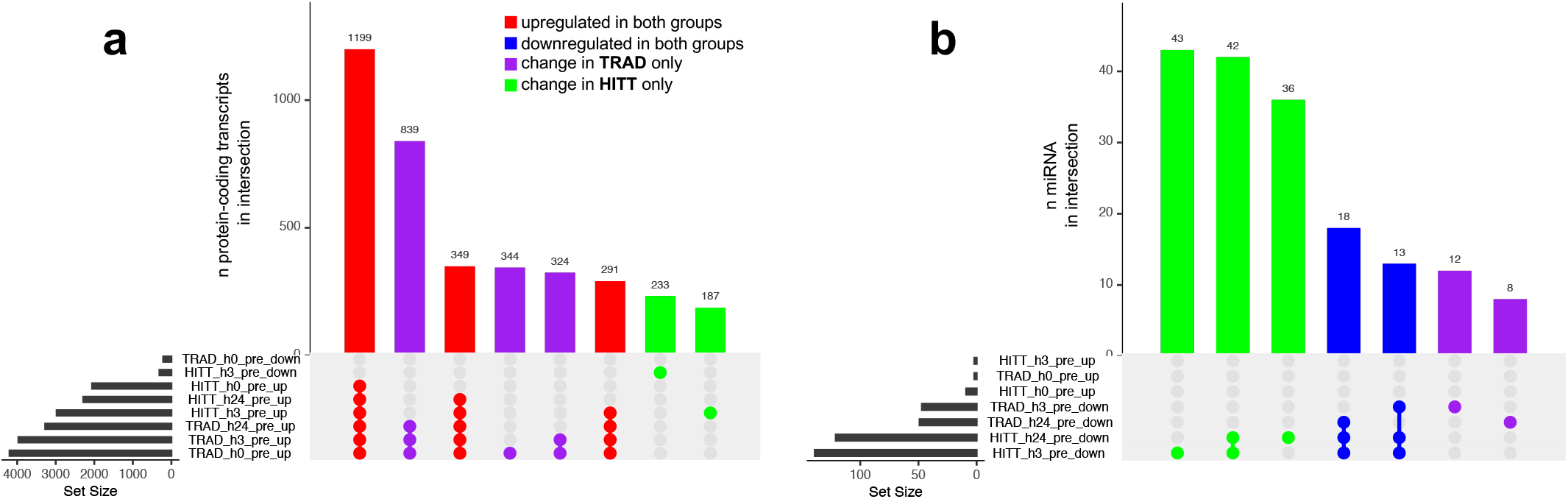
UpsetR plots indicating shared overlap (intersection) of directional transcript expression changes vs. pre-exercise in **(a)** protein-coding transcripts and **(b)** miRNA in serum extracellular vesicles (EVs). Bar height represents the number of genes in each intersection, whereas dots represent comparison[s] in which transcripts were significantly up- or down-regulated vs. pre-exercise (|log_2_FC| >1 and FDR<0.05). Gene intersections are shown in red where upregulation was detected at any time point independently of group, in blue where downregulation was detected independently of group, in purple where a change was detected in TRAD only, and in green where a change was detected in HITT only. Gene ontology analysis for each intersection in panel **a** is available in Supplementary Table 2.

#### Serum EV Small RNA

Following acute exposure to TRAD, 2 small transcripts were upregulated at h0 post-exercise (both miRNA), compared to 24 (38% miRNA) in HITT, including the 2 seen in TRAD. No small transcripts in serum EVs were significantly downregulated from pre-exercise in either group at this time point. At h3 post-exercise in TRAD, 4 small transcripts (0% miRNA) were upregulated, and 8 small transcripts (25% miRNAs) were upregulated in HITT, including the 4 significantly changed in TRAD. Conversely, 52 transcripts (90% miRNA) were downregulated in TRAD at h3, and 148 (94% miRNA) were downregulated in HITT. Of these, 32 (30 miRNA) were shared between groups. Finally, at h24 post-exercise, a total of 7 small transcripts (no miRNA) were upregulated in TRAD, and none were significantly changed in HITT. A total of 50 small transcripts (98% miRNA) were downregulated in TRAD compared to 126 (96% miRNA) in HITT. Of these, 29 (all miRNA) were common to both groups. Because this data stream exhibited a contrary pattern wherein HITT elicited a larger gene expression response, Supplementary Fig. 4 shows the patterns in miRNA at h0 (a), h3 (b), and h24 (c) post-exercise with respect to the FDR and log_2_FC vs. pre-exercise in HITT only, with filled shapes representing where a significant change in expression was also detected in TRAD. Figure 4b shows that the three largest intersections in small RNA from serum EVs were unique to HITT only.

#### Serum EV Circular RNA

At h0 post-exercise, 1 circRNA was upregulated and 0 downregulated in TRAD vs. 30 upregulated and 6 downregulated in HITT. No significant changes were seen in TRAD circRNA expression at h3 or h24 post-exercise. However, at h3 post-exercise, 2 circRNA were upregulated and 3 downregulated in HITT. At h24, 3 were upregulated and 1 downregulated in HITT. Given that the response in serum EV circular RNA was overall more muted compared to the other data streams, these acute changes are not visually presented.

### Exploratory Transcriptomic Network Discovery

#### Regulatory Networks in Skeletal Muscle

Skeletal muscle transcript expression networks were constructed from a total of 21,320 “responsive” transcripts that were different from pre-exercise at any time point based on FDR<0.20. PLIER-guided SVD generated 40 LVs from skeletal muscle transcripts, of which 23 showed a significant (p<0.00125) main effect of exercise (11 early, 6 late, 6 sustained). Table 5 shows the manually-selected biological annotations associated with each of the 23 LVs, derived from detailed results of GO/KEGG enrichment analysis available in Supplementary Table 3. LVs demonstrating a group x time interaction effect at the post-hoc level of significance are presented in Fig. 5a-e, along with the proportion of other RNA biotypes associated with the LV that passed 50% of the maximum loading threshold. A full list of all transcripts above this threshold is available for each of the 23 LVs in Supplementary Table 4. In particular, skeletal muscle transcript LVs 1 (intracellular signaling) and 34 (lipid metabolism) were significantly different at h3 post-exercise in both groups and also different at h3 between HITT vs. TRAD. LVs 4 (kinase activity, muscle tissue development) and 14 (negative regulation of muscle tissue development) were significantly different at h3 post-exercise in TRAD and different from HITT at the same time point. LV20 (skin and cardiac muscle development, hormone response) was different at h3 post-exercise in HITT and different from TRAD at h3; within TRAD, LV20 was different at h3 vs. h24.

**Table 5:**
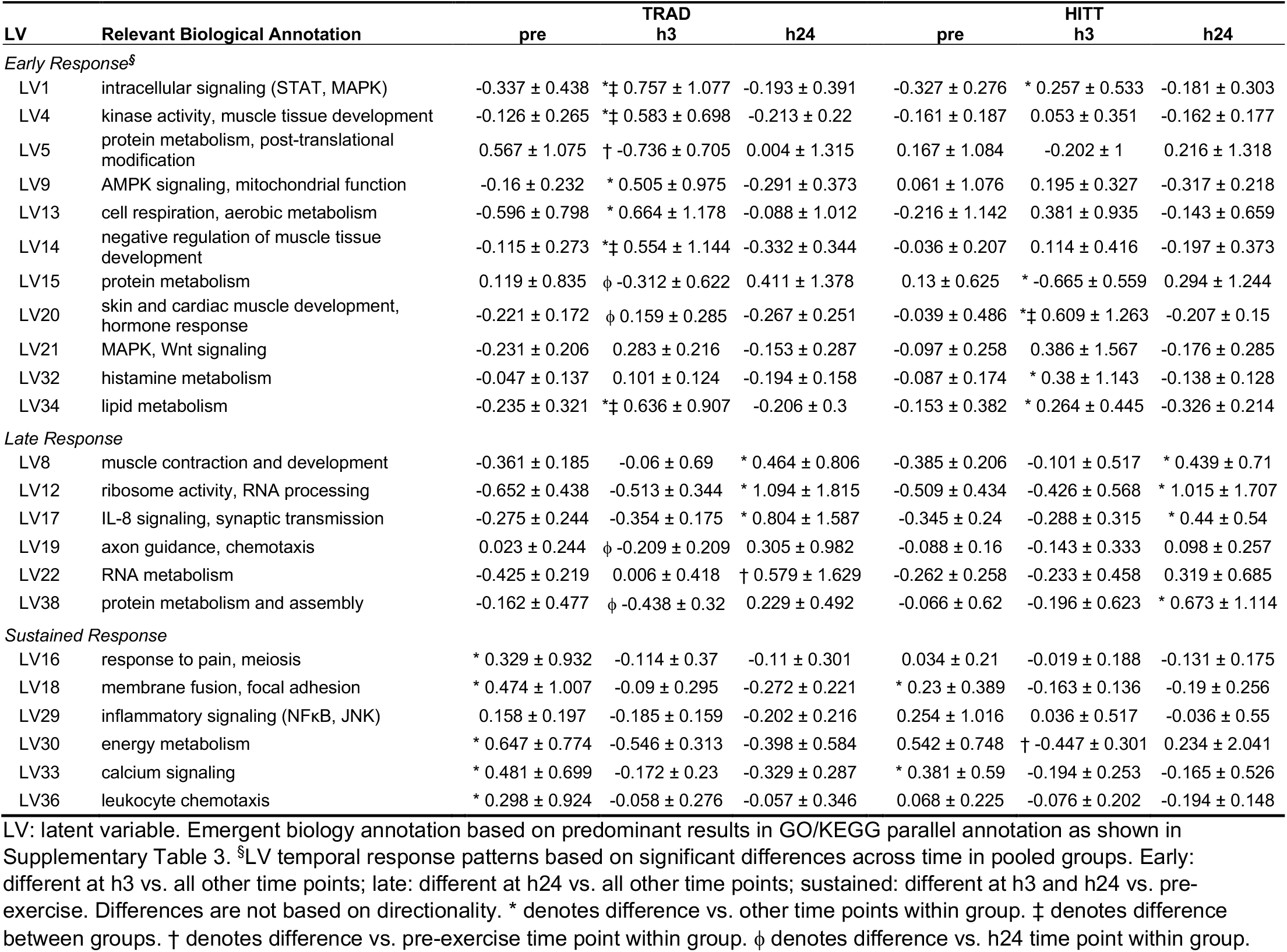
Values for LVs of interest in the muscle transcriptomics network analysis for each exercise dose.

**Fig. 5:**
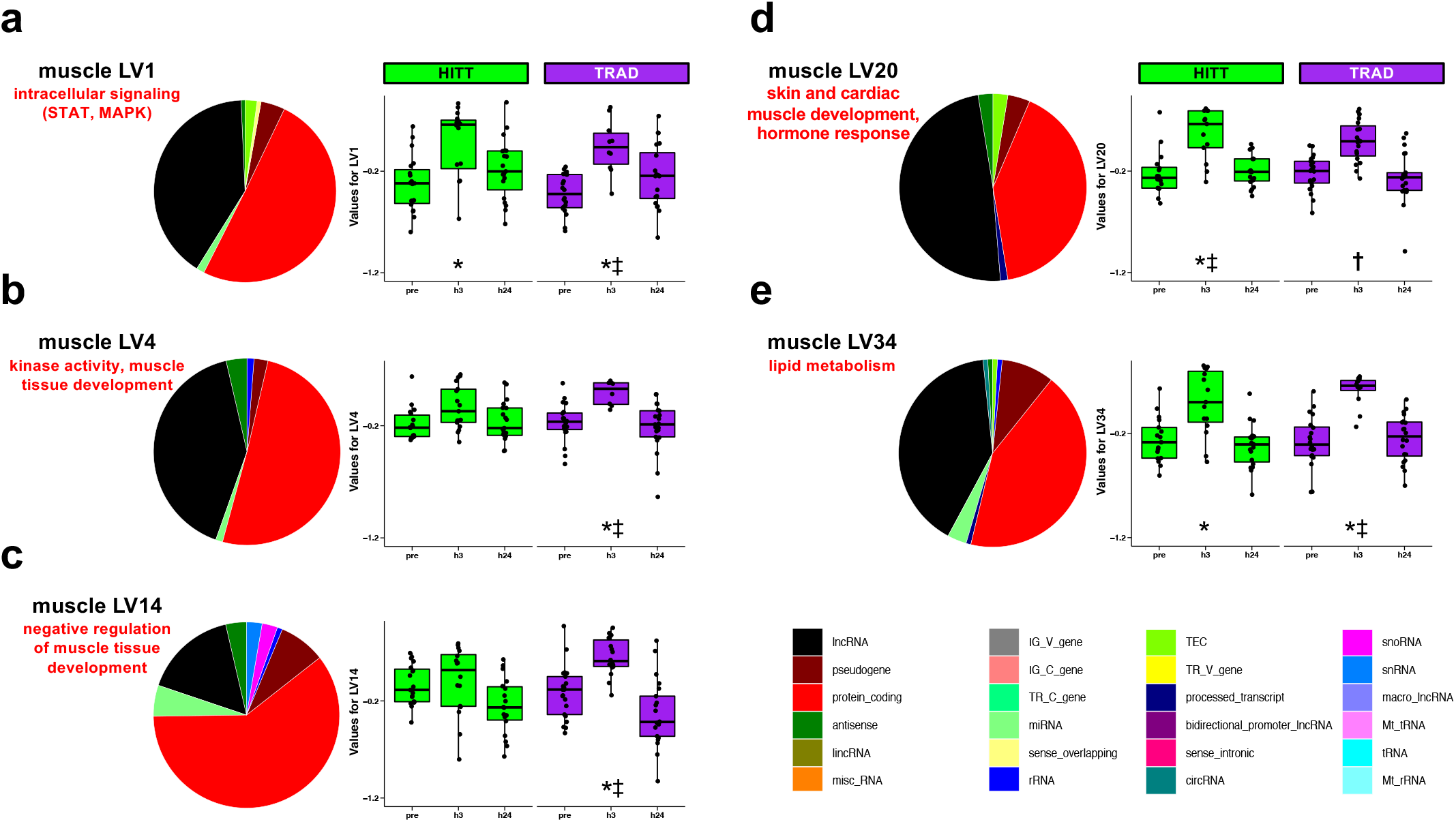
Latent variables (LVs) in skeletal muscle where a group x time difference in LV was detected via post-hoc testing of PLIER-generated LVs. For each LV (**a-e**), the fraction of transcripts most highly representative of the LV are summarized for biotype, and then the protein-coding fraction of each LV was annotated using a combination of Gene Ontology and KEGG, as summarized on the right of each pie chart. Additional LVs exhibited a main effect of exercise across both groups and are shown in Table 5. * denotes post-hoc p<0.05 vs. pre-exercise within group; ‡ denotes post-hoc p<0.05 vs. other group at the same time point; † denotes post-hoc p<0.05 vs. h24 time point within group.

#### Networks in Serum Extracellular Vesicles

Serum EV transcript expression networks were constructed from a total of 56,407 “responsive” transcripts that were different from pre-exercise at any time point based on FDR<0.20. PLIER-guided SVD generated 32 LVs from serum EV transcripts, of which 4 showed a significant (p<0.00156) main effect of exercise. Table 6 shows the general biological annotations associated with each of the four LVs, derived from detailed results of GO/KEGG analysis available in Supplementary Table 5. These four LVs are presented in Fig. 6a-d, along with the proportion of other RNA biotypes associated with the LV that passed 50% of the maximum loading threshold. A list of all transcripts above this threshold is available for each of the four LVs in Supplementary Table 6. Briefly, serum EV transcript LVs 10 (protein metabolism and biosynthesis), 17 (leukocyte activation and immune response), and 19 (blood coagulation and platelet biology) were significantly different at every post-exercise time point vs. pre-exercise in TRAD only. Serum EV LV2 (sensory perception and cell mobility) was different post-exercise overall but did not survive post-hoc tests when the groups were separated.

**Table 6:** Values for LVs of interest in the serum EV transcriptomics network analysis for each exercise dose.

| LV | Relevant Biological Annotation | TRAD |  |  |  | HITT |  |  |  |
| --- | --- | --- | --- | --- | --- | --- | --- | --- | --- |
|  |  | pre | h0 | h3 | h24 | pre | h0 | h3 | h24 |
| Sustained Response<sup>§</sup> |  |  |  |  |  |  |  |  |  |
| LV2 | sensory perception and cell mobility | -0.853 ± 0.773 | -0.224 ± 1.138 | 0.109 ± 1.192 | 0.061 ± 0.954 | 0.06 ± 2.201 | -0.186 ± 0.751 | 0.658 ± 1.183 | 0.793 ± 1.058 |
| LV10 | protein metabolism and biosynthesis | * 1.169 ± 2.963 | -0.312 ± 0.369 | -0.229 ± 0.536 | 0.011 ± 0.497 | 0.159 ± 0.868 | -0.348 ± 0.294 | -0.431 ± 0.242 | -0.326 ± 0.309 |
| LV17 | leukocyte activation and immune response | * 0.93 ± 2.296 | -0.248 ± 0.376 | -0.165 ± 0.634 | -0.001 ± 0.414 | 0.134 ± 0.704 | -0.265 ± 0.355 | -0.37 ± 0.19 | -0.262 ± 0.326 |
| LV19 | blood coagulation and platelet biology | * 0.842 ± 1.137 | -0.261 ± 0.409 | -0.336 ± 0.378 | -0.074 ± 0.467 | 0.935 ± 2.618 | -0.329 ± 0.288 | -0.532 ± 0.2 | -0.402 ± 0.297 |

LV: latent variable. Emergent biology annotation based on predominant results in GO/KEGG parallel annotation as shown in Supplementary Table 5. <sup>§</sup>LV temporal response patterns based on significant differences across time in pooled groups. Sustained: different at h0, h3, and h24 vs. pre-exercise (LV2 follows general pattern but was not statistically different at h0 vs. pre). Differences are not based on directionality. \* denotes significant difference vs. other time points within group.

**Fig. 6:**
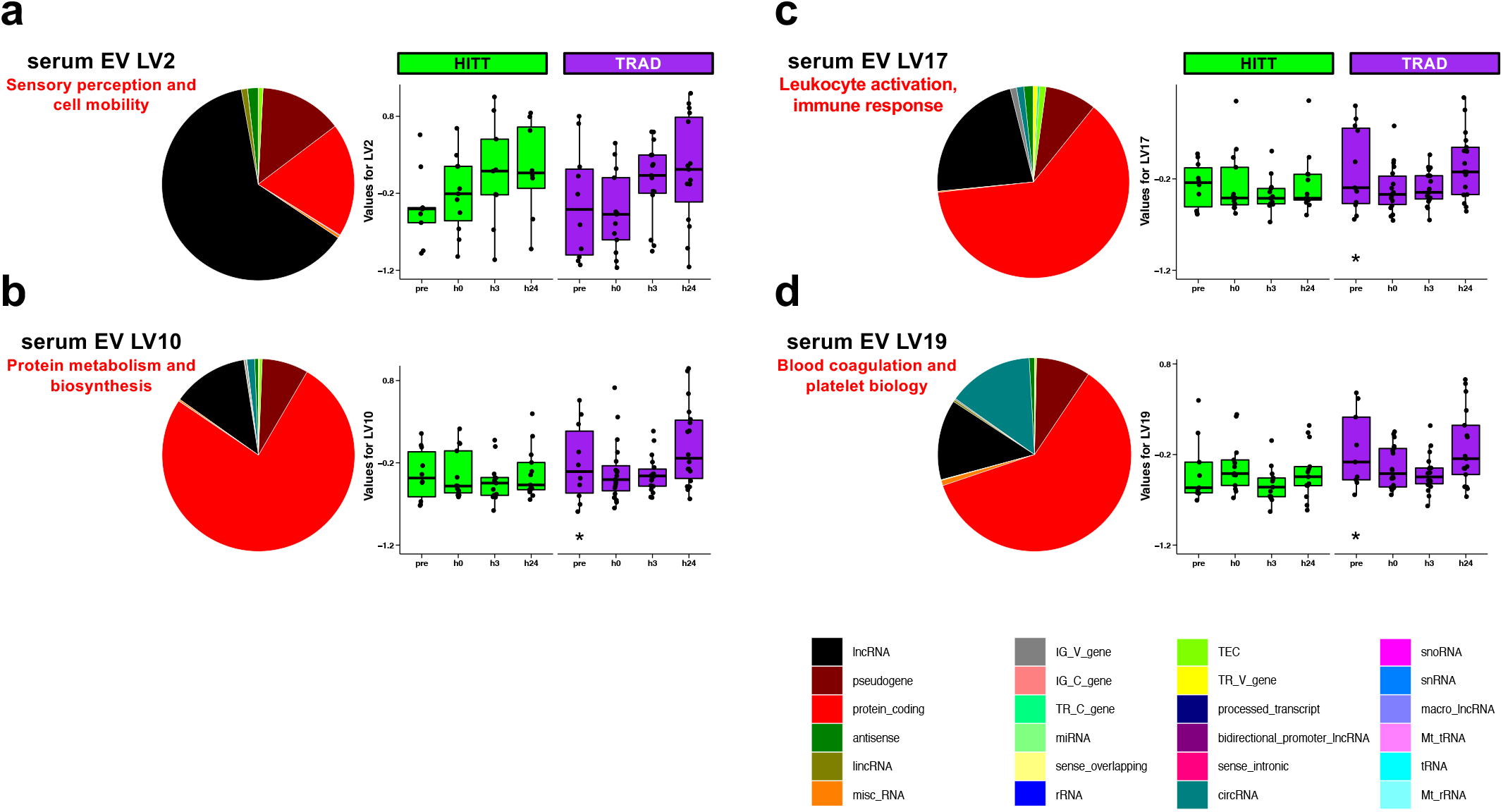
Latent variables (LVs) in serum extracellular vesicles (EVs) where a difference in LV was detected via post-hoc testing of PLIER-generated LVs. For each LV (**a-d**), the fraction of transcripts most highly representative of the LV are summarized for biotype, and then the protein-coding fraction of each LV was annotated using a combination of Gene Ontology and KEGG, as summarized on the right of each pie chart. * denotes post-hoc p<0.05 vs. pre-exercise within group.

## DISCUSSION

Exercise remains the most powerful medicine in promoting health and physical resilience to stress; thus, it is critically important to understand the mechanisms underlying these benefits. Acute exercise provides valuable insight into molecular patterns that may form the basis of adaptations conferred through repeated exposures over the long-term (i.e., consistent exercise training). This could aid in establishing direction toward personalized exercise training prescriptions that leverage inter-individual variation in molecular phenotype and associated whole-body phenotypes. In the present study, we compared the acute transcriptomic responses to two potent yet distinct, combined-exercise prescriptions and examined the interrelationships across transcript biotypes. Our findings provide insight into the temporal dynamics of the transcriptome circuitry and patterns that may suggest regulatory or other molecular interactions among transcripts. To our knowledge, this is the largest molecular mapping study of its kind to date in terms of sample size, comparisons across two randomized exercise doses, and depth and breadth of skeletal muscle and serum EV transcriptomic profiling (small, long, and circular RNA encompassing a total of 20 biotypes).

### Impact of Exercise Dose

In both the transcript-level and network-level analyses, the overall transcriptomic response was larger and/or more consistent in TRAD vs. HITT, although the composition of differentially-expressed transcripts in both serum EV and skeletal muscle (e.g., % protein-coding fraction) appeared generally similar between groups. While it may be argued that similar overall directional patterns were evident in HITT and declaration of significant difference was restricted by the rigorous yet still arbitrary significance and fold-change cutoffs (Fig. 2), this effect would likely be seen regardless of the specific thresholds imposed. Thus, it is potentially important that in well-balanced, similarly-sized groups with no apparent different physical characteristics at baseline, a larger response consistently emerged in TRAD. A potential explanation for these findings is that the TRAD protocol lasted approximately twice as long in duration as HITT, and this might account for lower variability across individuals by the time the biopsies and blood draws were collected. However, the more likely explanation is that the longer exercise regimen presented a sustained, unique metabolic challenge not experienced by participants undergoing HITT. Detection of truly divergent responses might require a larger sample size in order to account for inter-individual response heterogeneity in the exercise-related response ability to detect changes. Nevertheless, heterogeneity itself is important and may present opportunities for follow-up research, particularly as it relates to prescription of and adaptation to HITT across individuals.

On the contrary, miRNA in serum EVs exhibited a greater response following HITT exercise, with most of these downregulated (Fig. 4b). As shown in Supplementary Fig. 4, these differences do not tend to cluster below a particular fold-change cutoff, as was apparent in the other data streams. Typically, miRNA are thought to act as post-transcriptional silencers; thus, a downregulation might be expected to lead to heightened availability of transcripts for translation. While we did not assess protein-level changes in response to acute exercise, it would be informative to establish whether chronic exposure to HITT might differentially impact the abundance of critical proteins that modulate some adaptations.

A comparison to a tissue enrichment atlas developed by members of our team (1) showed a key miRNA downregulated in HITT only at h24 (hsa-mir-9-3) is exclusively and significantly enriched in central nervous system regions including the cerebellum, cerebral cortex, spinal cord, and brain white matter. This suggests a potential crosstalk originating in the nervous system that is unique to the high intensity stimulus, possibly necessitated by the nature of the bout (a timed sequence of varied functional exercises) and its demand on cognition, concentration, memory, neuromuscular control (balance, proprioception, coordination), maximal motor unit activation, and other neural processes. In support, other neuroactive molecules have been reported to exhibit intensity-dependent effects (8, 31), while accustomed moderate-intensity cycling facilitates reduced cognitive burden during exercise (27). It could be highly informative to establish whether this effect of HITT is sustained over time and has long-term benefits for cognition over TRAD, but this would likely be best illustrated in a population other than young, healthy adults with normal cognitive function.

### Insight from Serum EVs

EVs are produced by nearly every cell and are a convenient and accessible reflection of the exercise “secretome” (55). Understanding their source of origin may provide key insight into the systemic effects of exercise, which impacts nearly every physiological system (19) and stimulates tissue crosstalk. While a large portion of differentially expressed transcripts were unique to serum EVs, 401 transcripts across all RNA biotypes were significantly impacted at all measured timepoints within both body compartments (Fig. 7), indicating a coordinated and consistent physiological response to the exercise stimulus. In spite of this, protein-coding transcripts and validated functional targets of differentially expressed miRNA covered a wide and non-overlapping range of biological processes (Fig. 8), including cytokine signaling, kinase activity, and growth factor activity for the targets of differentially expressed miRNA, and G-protein coupled signaling, hormone activity, and ion channel regulation for differentially expressed protein-coding genes. This is perhaps not entirely surprising, given that miRNA act post-transcriptionally and would not necessarily be expected to correlate with changes in expression of their potential target genes. Additional inter-regulatory dynamics, such as tissue crosstalk and temporal regulation (e.g., miRNA changing in serum at post-exercise h0 impacting translation of mRNA in muscle at post-exercise h3 or h24) must be taken into account when attempting to integrate such multidimensional data. For these reasons, the agnostic SVD approach in PLIER provides a solid mathematical basis by which relatedness across transcript biotypes is uncovered. Continued mechanistic experiments to understand their physical interactions are certainly necessary to inform design of future multiomics studies.

**Fig. 7:**
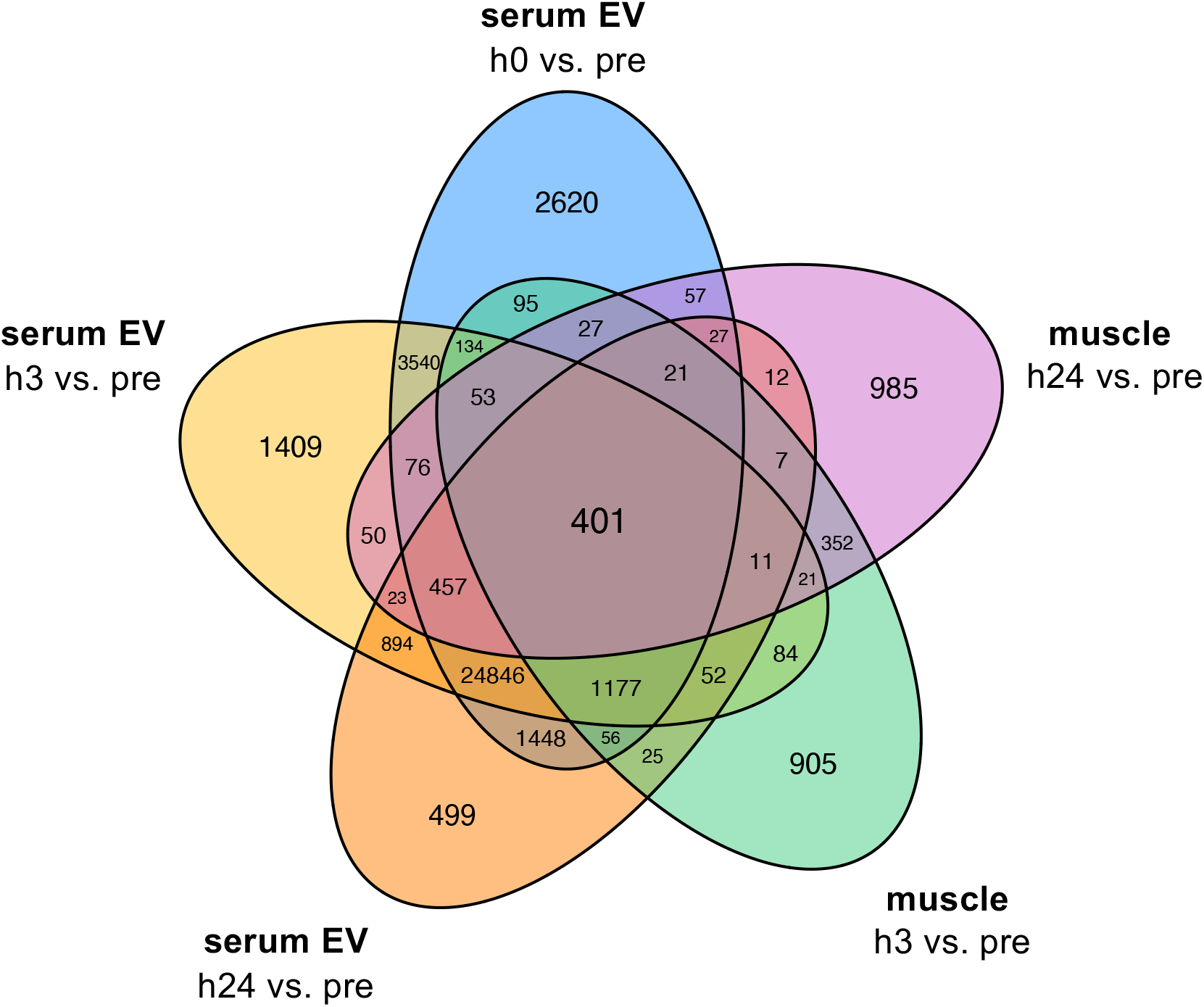
Venn diagram showing shared overlap of differentially expressed transcripts in both muscle and serum extracellular vesicles (EVs) at all post-exercise time points (0h, h3, and h24) vs. pre-exercise (pre). All transcripts satisfying both |log2FC| >1 and FDR<0.05 were collapsed regardless of directional change or RNA biotype.

**Fig. 8:**
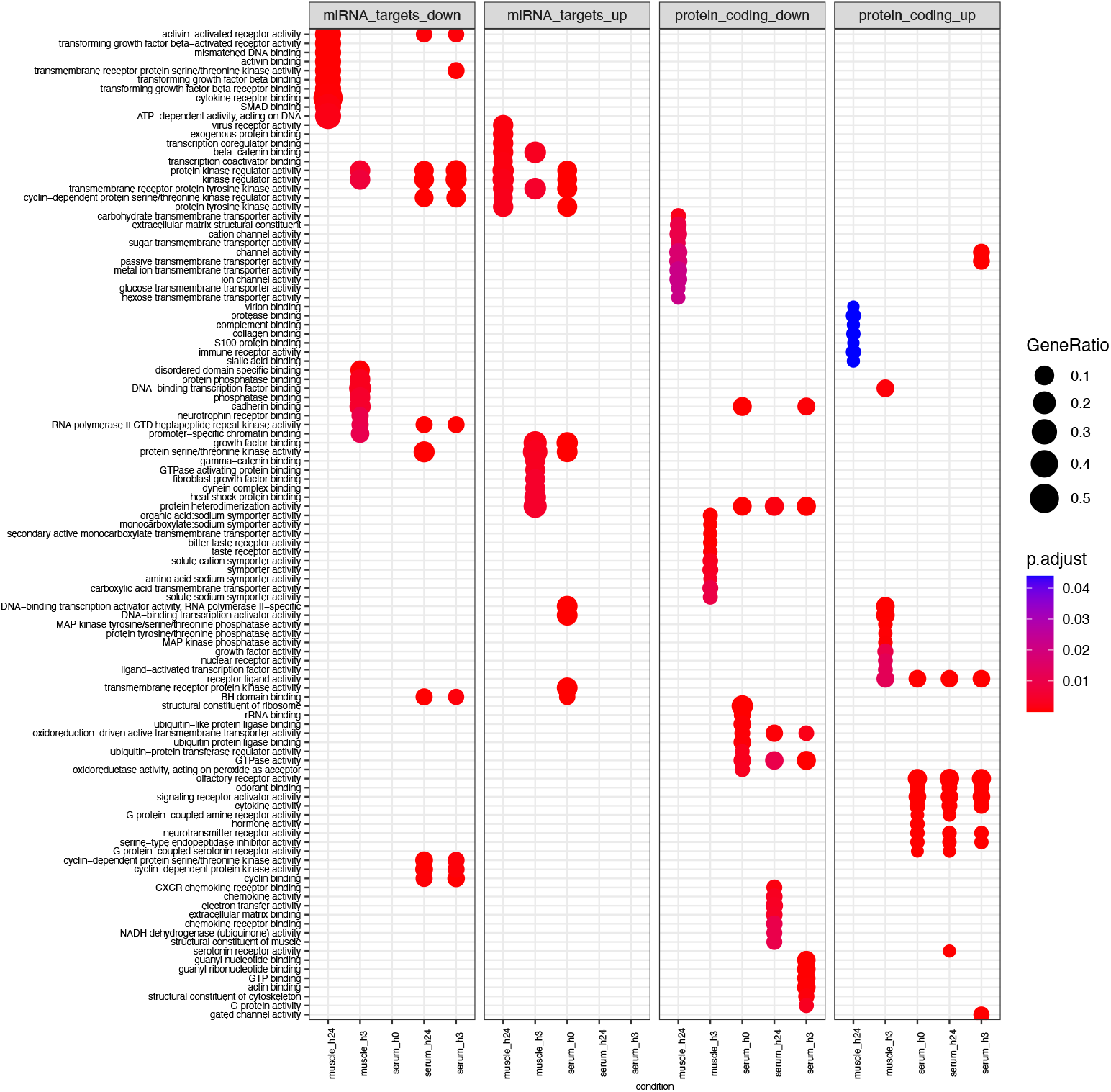
Biological processes associated with differentially expressed transcripts in skeletal muscle and serum extracellular vesicles at all post-exercise time points (h0, h3, and h24) with FDR < 0.05 and absolute log_2_fold-change >1 vs. pre-exercise, as curated by Gene Ontology. For miRNA targets (left two panels), multiMiR (v1.16) was used to identify the top 5% of predicted protein-coding mRNAs that functionally interact with miRNA differentially expressed following acute exercise. Targets were combined for all miRNA upregulated (up) or downregulated (down) at a given time point within muscle and serum extracellular vesicles and entered into gene ontology analysis. Differentially expressed protein-coding transcripts (right two panels) were directly input into gene ontology. GeneRatio (circle size) represents the proportion of genes shared between gene query and biological pathway, whereas likelihood of overlap based on the reference genome is represented by p.adjust (circle color).

As an exploratory approach to interpret patterns in the protein-coding transcriptome after acute HITT and TRAD exercise, we used the Human Protein Atlas (53) to map transcripts to protein expression patterns enriched at or above a medium level of confidence and supported reliability or better in bodily tissues of interest (Supplementary Fig. 5), revealing an integrative whole-body response to the acute exercise stimulus via shared genes upregulated at all timepoints in both doses. Skeletal muscle-enriched genes formed the basis of the largest intersection (n=18) and were mostly associated with processes of contractile function and muscle development. It could be meaningful to establish the source of these EVs and whether they may be originating from or targeting skeletal muscle. Tissue-specific EV patterns such as surface markers and/or cargo may be measured for this purpose in future investigations. The Human Protein Atlas also identified clusters of genes enriched in pancreas, stomach, and adipose, all tissues that are likely engaged in metabolic flux following the energy demands of acute exercise. The dramatically asymmetrical response seen in serum EV long RNA (strongly positive) was critically evaluated during QC and determined not to be the result of technical artifact (e.g., laboratory technician or sample batching). While the asymmetry persisted, it may be of interest that the imbalanced ratio of up- to downregulated transcripts in serum EVs after exercise is notably diminished when considering only the proportion of protein-coding genes, which was remarkably consistent at each post-exercise timepoint (∼12-14% of all differentially expressed transcripts). While the biological roles of the larger, non-coding fraction of differentially expressed transcripts is unclear, this imbalance may suggest an overall transcriptional stress response to acute exercise throughout the body. It would be informative to establish whether this signature persists in the trained state.

PLIER provided key insight into the potential relationship of upregulated transcripts to biological pathways associated with circulating blood cell types (naturally not in serum). For instance, serum EV LV17 was related to leukocyte activation and immunity, whereas serum EV LV19 was related to clotting and platelet biology. Supporting this as a likely source of variation, a recent study examining the multiomic patterns in a specialized subset of blood cells (e.g., peripheral blood mononuclear cells) found that the transcriptomic response following acute exercise is relatively well-balanced between the number of up- and downregulated genes(4). While insight into more direct contributions of different blood cell types might have been gained leveraging a prior knowledge database (29), we elected to perform PLIER agnostically in order to retain unannotated transcripts, which would otherwise be automatically removed by the algorithm (29). As such, a follow-up analysis would be necessary to establish whether the observed changes reflect the expected exercise-induced shifts in circulating cell composition (47), under the potentially tenuous assumption that resting gene expression signature is representative of the post-exercise state in these blood cells.

As opposed to other LVs of interest associated with mostly protein coding mRNA (>60% protein-coding in serum EVs, ∼46% in skeletal muscle), serum EV LV2 was unique in that it was associated with a large component of lncRNAs (63% in top 50^th^ percentile). This population of transcripts has been recently studied in the context of exercise adaptation (3, 6), although its biological roles are incompletely understood. The ten lncRNAs loaded into serum EV LV2 are poorly annotated, hinging interpretation of this LV completely on the relatively smaller protein-coding component (19%). However, future transcriptome-wide studies employing PLIER may establish whether these lncRNAs and other under-annotated biotypes consistently associate with the same functional categories of protein-coding genes. Thus, it may be possible computationally and iteratively to build a resource of likely biological roles for these biotypes to be experimentally validated via standard biochemical assays.

### Temporal Dynamics of the Transcriptomic Response

Temporally classifying the transcript LVs reveals the overall dynamics in response to the two divergent exercise bouts. Of 23 LVs in muscle that showed a main effect of exercise, 11 were responsive at the early time point (i.e., significantly different from both other time points at h3 only). These played diverse roles including activation of multiple cellular signaling cascades implicated in energy metabolism (AMPK), growth (Wnt) and inflammation (e.g., MAPK, STAT). Considerable information could be gained by exploring the non-protein-coding fraction of these LVs and potential regulatory roles of transcripts in a cell/tissue culture model or other laboratory-based assay.

Previous studies have established 3-4h post-exercise as a highly transcriptionally active time point (26). While the present findings substantiate this, the temporal gene expression response within skeletal muscle tissue generally suggests that the molecular environment is still perturbed from rest at h24 after exposure to the exercise bout. In support, when PLIER was used to collapse transcripts into LVs, 12 of 23 that exhibited a significant time effect were different at h24 (i.e., either late or sustained). The “late” LVs (different at h24 only) in skeletal muscle were primarily associated with protein-coding genes linked to processes related to protein synthesis and metabolism, chemokine signaling, and neuromuscular communication. This suggests a role for these transcripts in recovery and mechanistic changes potentially underlying gains in strength, power, and muscle mass.

In serum EVs, all four LVs that showed a main effect of exercise were sustained at h24. This may be due to the unaccustomed nature of the exercise bout itself in untrained individuals. While all participants underwent four progressive familiarization sessions to reduce undue muscular pain and soreness, they remained relatively exercise-naïve. We suspect some of the molecular responses sustained at h24 may not be as protracted in exercise-trained individuals. It is well-established that a single exposure to an unaccustomed bout can reduce the magnitude of the effect of subsequent exposures, particularly in terms of a classical inflammatory response (15). Fittingly, transcripts associated with serum EV LV17 were linked to processes including leukocyte activation and immune response. While acute inflammation is considered important for exercise-induced muscle adaptation (21, 26, 39), chronic elevation has negative consequences. As a comparator, testing for these sustained response muscle LVs in trained participants would perhaps be a valuable means of filtering LVs (and identifying novel member genes) most likely to promote positive exercise adaptations. Studying the response of transcripts associated with LV17 to a chronic training regimen might provide such insight. Additionally, while typically restricted to skeletal muscle, prolonged pro-inflammation often seen before (51) and after (21) training in older adults is thought to be a barrier to adaptation. It could be informative to establish whether similar patterns in LV17 are upheld after unaccustomed exercise in older populations.

### Future Directions

While this study did not investigate the influence of sex, both males and females were included in relative balance by design (randomization stratified by sex). Thus, we did not seek to mathematically “erase” the influence of sex, which is thought to be an inappropriate practice in sex-cognizant research (34). Nevertheless, it is possible to leverage these data sets toward an investigation of sex, albeit somewhat restricted by the small sample size (n=∼10 per sex per exercise dose), using tools such as PLIER and PCA to reduce data dimensionality.

Additional investigations in populations of different age, health, or training status will also likely benefit from this in-depth inclusive description of short-term transcriptomic changes following exposure to an acute exercise stimulus. By continuing to build an extensive repository of rigorous data sets across molecular levels and by enriching the computational tools available to integrate and interpret them, a clearer understanding of the normal, healthy transcriptome map of exercise may eventually be built. By using this resource as a comparator data set, it may become possible to identify and target biological pathways dysregulated by disease in order to maximize exercise responsiveness for the greatest number of individuals (18) or augment the impact of combinatorial therapies for disease management.

Given the complex and integrated signaling responses to an acute exercise stress, examining exercise-induced changes at multiple molecular levels (i.e., via multiomics) is likely to provide even further insight into coordination and control of molecular processes underlying adaptation to exercise. This goal has led to the recent establishment of noteworthy efforts such as the Molecular Transducers of Physical Activity Consortium (MoTrPAC) (44), the Athlome (46), and numerous smaller-scale studies (37) designed to characterize molecular responses to exercise. While the larger studies remain years away from interpretation and distribution of meaningful findings, they are likely to provide valuable multiomic data that may be leveraged as a healthy comparator data set for independent examinations of molecular phenotypes across levels. The present findings would be similarly expanded by integrating other ‘omics (e.g., metabolomics, proteomics, methylomics), assessing alternative post-transcriptional splicing, correlating patterns with phenotypes (e.g., any measures shown in Table 1) across individuals to predict or explain phenotypic or response heterogeneity (18), and characterizing the molecular responses to chronic HITT and TRAD exercise training (30).

## CONCLUSION

While both traditional and high-intensity exercise confer numerous health- and performance-related benefits when completed chronically, the underlying mechanisms driving these adaptations have not been directly compared. Our findings suggest that, at the selected time points, traditional exercise elicits a stronger or more consistent transcriptome-wide response in skeletal muscle and serum EVs of young, healthy adults. Additionally, our findings point to patterns across transcripts that may aid in laying the foundation for annotation of transcripts based on parameters other than the cellular behavior of functional proteins. Continued attention should be directed towards these under-studied biotypes, especially since they exhibit considerable plasticity following a bout of exercise and are therefore likely to play roles in molecular regulation and long-term adaptation. When integrated in a multiomic fashion with other molecular responses to acute and chronic exercise, the transcriptome may provide guidance towards optimizing exercise prescription and maximizing the benefits while increasing enjoyment and reducing time burden for exercise participants. This is of utmost importance if exercise is to become a public health focus and a habitual component of a lifestyle change for individuals across the lifespan.

## ACKNOWLEDGMENTS

The authors thank all participants in this trial, in addition to the study coordinators, interventionists, all involved with physical performance testing, biospecimen collection, and inventorying, and the UAB Center for Exercise Medicine Core Muscle Research Lab and Exercise Clinical Training Facility personnel. Parent randomized clinical trial supported by DoD ONR N000141613159. ZAG is supported by the Dept. of Veterans Affairs Rehabilitation Research & Development Service (IK2RX002781). Research reported in this publication included work performed in the Integrated Mass Spectrometry Shared Resource supported by the National Cancer Institute of the National Institutes of Health under grant P30CA033572.

## AUTHOR CONTRIBUTIONS

- KML, KJ, MJH, PP, TJB, MMB conceived and designed research
- KML, SMO, JSM, DD, MBB, CJK, BP, SCT, RSS, RR, EA, EH, JA, AB, BM, JP, JST, IA, KP performed experiments
- KML, JSM, MEL, EA, EH, JA, JP analyzed data
- KML, ZAG, JSM, SMO, DD, KJ, KJ, MJH, PP, TB, MMB interpreted results of experiments
- KML drafted the manuscript
- KML prepared code and figures
- KML, ZAG, JSM, SMO, DD, KJ, MJH, PP, RR, EA, EH, TJB, MMB edited and revised manuscript
- KML, ZAG, JSM, SMO, DD, MBB, CJK, MEL, BP, SCT, RSS, KJ, MJH, PP, RR, EA, EH, JA, AB, BM, JP, JST, AS, IA, KP, TJB, MMB approved final version of manuscript

## REFERENCES

1. Alsop E, Meechoovet B, Kitchen R, Sweeney T, Beach TG, Serrano GE, Hutchins E, Ghiran I, Reiman R, Syring M, Hsieh M, Courtright-Lim A, Valkov N, Whitsett TG, Rakela J, Pockros P, Rozowsky J, Gallego J, Huentelman MJ, Shah R, Nakaji P, Kalani MYS, Laurent L, Das S, Van Keuren-Jensen K. A Novel Tissue Atlas and Online Tool for the Interrogation of Small RNA Expression in Human Tissues and Biofluids. Front Cell Dev Biol. 10: 804164, 2022.

2. Bar-Or O. The Wingate anaerobic test. An update on methodology, reliability and validity. Sports Med. 4: 381–394, 1987.

3. Bonilauri B, Dallagiovanna B. Long Non-coding RNAs Are Differentially Expressed After Different Exercise Training Programs. Front Physiol. 11: 567614, 2020.

4. Contrepois K, Wu S, Moneghetti KJ, Hornburg D, Ahadi S, Tsai MS, Metwally AA, Wei E, Lee-McMullen B, Quijada JV, Chen S, Christle JW, Ellenberger M, Balliu B, Taylor S, Durrant MG, Knowles DA, Choudhry H, Ashland M, Bahmani A, Enslen B, Amsallem M, Kobayashi Y, Avina M, Perelman D, Schussler-Fiorenza Rose SM, Zhou W, Ashley EA, Montgomery SB, Chaib H, Haddad F, Snyder MP. Molecular Choreography of Acute Exercise. Cell. 181: 1112–1130 e1116, 2020.

5. Conway JR, Lex A, Gehlenborg N. UpSetR: an R package for the visualization of intersecting sets and their properties. Bioinformatics. 33: 2938–2940, 2017.

6. De Sanctis P, Filardo G, Abruzzo PM, Astolfi A, Bolotta A, Indio V, Di Martino A, Hofer C, Kern H, Lofler S, Marcacci M, Marini M, Zampieri S, Zucchini C. Non-Coding RNAs in the Transcriptional Network That Differentiates Skeletal Muscles of Sedentary from Long-Term Endurance-and Resistance-Trained Elderly. Int J Mol Sci. 22: 2021.

7. Dobin A, Davis CA, Schlesinger F, Drenkow J, Zaleski C, Jha S, Batut P, Chaisson M, Gingeras TR. STAR: ultrafast universal RNA-seq aligner. Bioinformatics. 29: 15–21, 2013.

8. Eaton M, Granata C, Barry J, Safdar A, Bishop D, Little JP. Impact of a single bout of high-intensity interval exercise and short-term interval training on interleukin-6, FNDC5, and METRNL mRNA expression in human skeletal muscle. J Sport Health Sci. 7: 191–196, 2018.

9. Estrada AL, Valenti ZJ, Hehn G, Amorese AJ, Williams NS, Balestrieri NP, Deighan C, Allen CP, Spangenburg EE, Kruh-Garcia NA, Lark DS. Extracellular vesicle secretion is tissue-dependent ex vivo and skeletal muscle myofiber extracellular vesicles reach the circulation in vivo. Am J Physiol Cell Physiol. 322: C246–C259, 2022.

10. Garner RT, Solfest JS, Nie Y, Kuang S, Stout J, Gavin TP. Multivesicular body and exosome pathway responses to acute exercise. Exp Physiol. 105: 511–521, 2020.

11. Giraldez MD, Spengler RM, Etheridge A, Godoy PM, Barczak AJ, Srinivasan S, De Hoff PL, Tanriverdi K, Courtright A, Lu S, Khoory J, Rubio R, Baxter D, Driedonks TAP, Buermans HPJ, Nolte-’t Hoen ENM, Jiang H, Wang K, Ghiran I, Wang YE, Van Keuren-Jensen K, Freedman JE, Woodruff PG, Laurent LC, Erle DJ, Galas DJ, Tewari M. Comprehensive multi-center assessment of small RNA-seq methods for quantitative miRNA profiling. Nat Biotechnol. 36: 746–757, 2018.

12. Greco S, Cardinali B, Falcone G, Martelli F. Circular RNAs in Muscle Function and Disease. Int J Mol Sci. 19: 2018.

13. Haddock CK, Poston WS, Heinrich KM, Jahnke SA, Jitnarin N. The Benefits of High-Intensity Functional Training Fitness Programs for Military Personnel. Mil Med. 181: e1508–e1514, 2016.

14. Hutchins E, Reiman R, Winarta J, Beecroft T, Richholt R, De Both M, Shahbander K, Carlson E, Janss A, Siniard A, Balak C, Bruhns R, Whitsett TG, McCoy R, Anastasi M, Allen A, Churas B, Huentelman M, Van Keuren-Jensen K. Extracellular circular RNA profiles in plasma and urine of healthy, male college athletes. Sci Data. 8: 276, 2021.

15. Hyldahl RD, Chen TC, Nosaka K. Mechanisms and Mediators of the Skeletal Muscle Repeated Bout Effect. Exerc Sport Sci Rev. 45: 24–33, 2017.

16. Kanehisa M, Sato Y, Kawashima M, Furumichi M, Tanabe M. KEGG as a reference resource for gene and protein annotation. Nucleic Acids Res. 44: D457–462, 2016.

17. Kelly NA, Ford MP, Standaert DG, Watts RL, Bickel CS, Moellering DR, Tuggle SC, Williams JY, Lieb L, Windham ST, Bamman MM. Novel, high-intensity exercise prescription improves muscle mass, mitochondrial function, and physical capacity in individuals with Parkinson’s disease. J Appl Physiol (1985). 116: 582–592, 2014.

18. Lavin KM, Bell MB, McAdam JS, Peck BD, Walton RG, Windham ST, Tuggle SC, Long DE, Kern PA, Peterson CA, Bamman MM. Muscle transcriptional networks linked to resistance exercise training hypertrophic response heterogeneity. Physiol Genomics. 53: 206–221, 2021.

19. Lavin KM, Coen PM, Baptista LC, Bell MB, Drummer D, Harper SA, Lixandrao ME, McAdam JS, O’Bryan SM, Ramos S, Roberts LM, Vega RB, Goodpaster BH, Bamman MM, Buford TW. State of Knowledge on Molecular Adaptations to Exercise in Humans: Historical Perspectives and Future Directions. Compr Physiol. 12: 3193–3279, 2022.

20. Lavin KM, Ge Y, Sealfon SC, Nair VD, Wilk K, McAdam JS, Windham ST, Kumar PL, McDonald MN, Bamman MM. Rehabilitative Impact of Exercise Training on Human Skeletal Muscle Transcriptional Programs in Parkinson’s Disease. Front Physiol. 11: 653, 2020.

21. Lavin KM, Perkins RK, Jemiolo B, Raue U, Trappe SW, Trappe TA. Effects of aging and lifelong aerobic exercise on basal and exercise-induced inflammation. J Appl Physiol (1985). 128: 87–99, 2020.

22. Lavin KM, Roberts BM, Fry CS, Moro T, Rasmussen BB, Bamman MM. The Importance of Resistance Exercise Training to Combat Neuromuscular Aging. Physiology (Bethesda). 34: 112–122, 2019.

23. Law CW, Alhamdoosh M, Su S, Dong X, Tian L, Smyth GK, Ritchie ME. RNA-seq analysis is easy as 1-2-3 with limma, Glimma and edgeR. F1000Res. 5: 2016.

24. Law CW, Chen Y, Shi W, Smyth GK. voom: Precision weights unlock linear model analysis tools for RNA-seq read counts. Genome Biol. 15: R29, 2014.

25. Law CW, Zeglinski K, Dong X, Alhamdoosh M, Smyth GK, Ritchie ME. A guide to creating design matrices for gene expression experiments. F1000Res. 9: 1444, 2020.

26. Louis E, Raue U, Yang Y, Jemiolo B, Trappe S. Time course of proteolytic, cytokine, and myostatin gene expression after acute exercise in human skeletal muscle. J Appl Physiol (1985). 103: 1744–1751, 2007.

27. Ludyga S, Gronwald T, Hottenrott K. The Athlete’s Brain: Cross-Sectional Evidence for Neural Efficiency during Cycling Exercise. Neural Plast. 2016: 4583674, 2016.

28. MacInnis MJ, Gibala MJ. Physiological adaptations to interval training and the role of exercise intensity. J Physiol. 595: 2915–2930, 2017.

29. Mao W, Zaslavsky E, Hartmann BM, Sealfon SC, Chikina M. Pathway-level information extractor (PLIER) for gene expression data. Nat Methods. 16: 607–610, 2019.

30. McAdam JS, et al. (Placeholder) Precision High-Intensity Training in Epigenetics: The PHITE Trial. TBA. 2022.

31. McKay BR, Nederveen JP, Fortino SA, Snijders T, Joanisse S, Kumbhare DA, Parise G. Brain-derived neurotrophic factor is associated with human muscle satellite cell differentiation in response to muscle-damaging exercise. Appl Physiol Nutr Metab. 45: 581–590, 2020.

32. Nair VD, Ge Y, Li S, Pincas H, Jain N, Seenarine N, Amper MAS, Goodpaster BH, Walsh MJ, Coen PM, Sealfon SC. Sedentary and Trained Older Men Have Distinct Circulating Exosomal microRNA Profiles at Baseline and in Response to Acute Exercise. Front Physiol. 11: 605, 2020.

33. Neufer PD, Dohm GL. Exercise induces a transient increase in transcription of the GLUT-4 gene in skeletal muscle. Am J Physiol. 265: C1597–1603, 1993.

34. O’Bryan SM, Connor KR, Drummer D, Lavin KM, Bamman MM. Considerations for Sex-Cognizant Research in Exercise Biology and Medicine. Front. Sports Act. Living. 2022.

35. Patro R, Duggal G, Love MI, Irizarry RA, Kingsford C. Salmon provides fast and bias-aware quantification of transcript expression. Nat Methods. 14: 417–419, 2017.

36. Perry CG, Lally J, Holloway GP, Heigenhauser GJ, Bonen A, Spriet LL. Repeated transient mRNA bursts precede increases in transcriptional and mitochondrial proteins during training in human skeletal muscle. J Physiol. 588: 4795–4810, 2010.

37. Pillon NJ, Gabriel BM, Dollet L, Smith JAB, Sardon Puig L, Botella J, Bishop DJ, Krook A, Zierath JR. Transcriptomic profiling of skeletal muscle adaptations to exercise and inactivity. Nat Commun. 11: 470, 2020.

38. Pine PS, Lund SP, Parsons JR, Vang LK, Mahabal AA, Cinquini L, Kelly SC, Kincaid H, Crichton DJ, Spira A, Liu G, Gower AC, Pass HI, Goparaju C, Dubinett SM, Krysan K, Stass SA, Kukuruga D, Van Keuren-Jensen K, Courtright-Lim A, Thompson KL, Rosenzweig BA, Sorbara L, Srivastava S, Salit ML. Summarizing performance for genome scale measurement of miRNA: reference samples and metrics. BMC Genomics. 19: 180, 2018.

39. Raue U, Jemiolo B, Yang Y, Trappe S. TWEAK-Fn14 pathway activation after exercise in human skeletal muscle: insights from two exercise modes and a time course investigation. J Appl Physiol (1985). 118: 569–578, 2015.

40. Roberts BM, Lavin KM, Many GM, Thalacker-Mercer A, Merritt EK, Bickel CS, Mayhew DL, Tuggle SC, Cross JM, Kosek DJ, Petrella JK, Brown CJ, Hunter GR, Windham ST, Allman RM, Bamman MM. Human neuromuscular aging: Sex differences revealed at the myocellular level. Exp Gerontol. 106: 116–124, 2018.

41. Rodosthenous RS, Hutchins E, Reiman R, Yeri AS, Srinivasan S, Whitsett TG, Ghiran I, Silverman MG, Laurent LC, Van Keuren-Jensen K, Das S. Profiling Extracellular Long RNA Transcriptome in Human Plasma and Extracellular Vesicles for Biomarker Discovery. iScience. 23: 101182, 2020.

42. Rozowsky J, Kitchen RR, Park JJ, Galeev TR, Diao J, Warrell J, Thistlethwaite W, Subramanian SL, Milosavljevic A, Gerstein M. exceRpt: A Comprehensive Analytic Platform for Extracellular RNA Profiling. Cell Syst. 8: 352–357 e353, 2019.

43. Rubenstein AB, Hinkley JM, Nair VD, Nudelman G, Standley RA, Yi F, Yu G, Trappe TA, Bamman MM, Trappe SW, Sparks LM, Goodpaster BH, Vega RB, Sealfon SC, Zaslavsky E, Coen PM. Skeletal muscle transcriptome response to a bout of endurance exercise in physically active and sedentary older adults. Am J Physiol Endocrinol Metab. 322: E260–E277, 2022.

44. Sanford JA, Nogiec CD, Lindholm ME, Adkins JN, Amar D, Dasari S, Drugan JK, Fernandez FM, Radom-Aizik S, Schenk S, Snyder MP, Tracy RP, Vanderboom P, Trappe S, Walsh MJ, Molecular Transducers of Physical Activity C. Molecular Transducers of Physical Activity Consortium (MoTrPAC): Mapping the Dynamic Responses to Exercise. Cell. 181: 1464–1474, 2020.

45. Saugstad JA, Lusardi TA, Van Keuren-Jensen KR, Phillips JI, Lind B, Harrington CA, McFarland TJ, Courtright AL, Reiman RA, Yeri AS, Kalani MYS, Adelson PD, Arango J, Nolan JP, Duggan E, Messer K, Akers JC, Galasko DR, Quinn JF, Carter BS, Hochberg FH. Analysis of extracellular RNA in cerebrospinal fluid. J Extracell Vesicles. 6: 1317577, 2017.

46. Sellami M, Elrayess MA, Puce L, Bragazzi NL. Molecular Big Data in Sports Sciences: State-of-Art and Future Prospects of OMICS-Based Sports Sciences. Front Mol Biosci. 8: 815410, 2021.

47. Shinkai S, Shore S, Shek PN, Shephard RJ. Acute exercise and immune function. Relationship between lymphocyte activity and changes in subset counts. Int J Sports Med. 13: 452–461, 1992.

48. Srinivasan S, Yeri A, Cheah PS, Chung A, Danielson K, De Hoff P, Filant J, Laurent CD, Laurent LD, Magee R, Moeller C, Murthy VL, Nejad P, Paul A, Rigoutsos I, Rodosthenous R, Shah RV, Simonson B, To C, Wong D, Yan IK, Zhang X, Balaj L, Breakefield XO, Daaboul G, Gandhi R, Lapidus J, Londin E, Patel T, Raffai RL, Sood AK, Alexander RP, Das S, Laurent LC. Small RNA Sequencing across Diverse Biofluids Identifies Optimal Methods for exRNA Isolation. Cell. 177: 446–462 e416, 2019.

49. Stec MJ, Thalacker-Mercer A, Mayhew DL, Kelly NA, Tuggle SC, Merritt EK, Brown CJ, Windham ST, Dell’Italia LJ, Bickel CS, Roberts BM, Vaughn KM, Isakova-Donahue I, Many GM, Bamman MM. Randomized, four-arm, dose-response clinical trial to optimize resistance exercise training for older adults with age-related muscle atrophy. Exp Gerontol. 99: 98–109, 2017.

50. Stensvold D, Viken H, Steinshamn SL, Dalen H, Stoylen A, Loennechen JP, Reitlo LS, Zisko N, Baekkerud FH, Tari AR, Sandbakk SB, Carlsen T, Ingebrigtsen JE, Lydersen S, Mattsson E, Anderssen SA, Fiatarone Singh MA, Coombes JS, Skogvoll E, Vatten LJ, Helbostad JL, Rognmo O, Wisloff U. Effect of exercise training for five years on all cause mortality in older adults-the Generation 100 study: randomised controlled trial. BMJ. 371: m3485, 2020.

51. Thalacker-Mercer A, Stec M, Cui X, Cross J, Windham S, Bamman M. Cluster analysis reveals differential transcript profiles associated with resistance training-induced human skeletal muscle hypertrophy. Physiol Genomics. 45: 499–507, 2013.

52. The Gene Ontology C. The Gene Ontology Resource: 20 years and still GOing strong. Nucleic Acids Res. 47: D330–D338, 2019.

53. Uhlen M, Fagerberg L, Hallstrom BM, Lindskog C, Oksvold P, Mardinoglu A, Sivertsson A, Kampf C, Sjostedt E, Asplund A, Olsson I, Edlund K, Lundberg E, Navani S, Szigyarto CA, Odeberg J, Djureinovic D, Takanen JO, Hober S, Alm T, Edqvist PH, Berling H, Tegel H, Mulder J, Rockberg J, Nilsson P, Schwenk JM, Hamsten M, von Feilitzen K, Forsberg M, Persson L, Johansson F, Zwahlen M, von Heijne G, Nielsen J, Ponten F. Proteomics. Tissue-based map of the human proteome. Science. 347: 1260419, 2015.

54. Volders PJ, Helsens K, Wang X, Menten B, Martens L, Gevaert K, Vandesompele J, Mestdagh P. LNCipedia: a database for annotated human lncRNA transcript sequences and structures. Nucleic Acids Res. 41: D246–251, 2013.

55. Whitham M, Febbraio MA. Redefining Tissue Crosstalk via Shotgun Proteomic Analyses of Plasma Extracellular Vesicles. Proteomics. 19: e1800154, 2019.

56. Winter J, Jung S, Keller S, Gregory RI, Diederichs S. Many roads to maturity: microRNA biogenesis pathways and their regulation. Nat Cell Biol. 11: 228–234, 2009.

57. Yeri A, Courtright A, Danielson K, Hutchins E, Alsop E, Carlson E, Hsieh M, Ziegler O, Das A, Shah RV, Rozowsky J, Das S, Van Keuren-Jensen K. Evaluation of commercially available small RNASeq library preparation kits using low input RNA. BMC Genomics. 19: 331, 2018.

58. Zhang Y, Parmigiani G, Johnson WE. ComBat-seq: batch effect adjustment for RNA-seq count data. NAR Genom Bioinform. 2: lqaa078, 2020.

