## Supplementary Fig. 1 for "Dynamic Transcriptomic Network Responses to Divergent Acute Exercise Challenges in Young Adults"

### SUPPLEMENTARY FIGURES

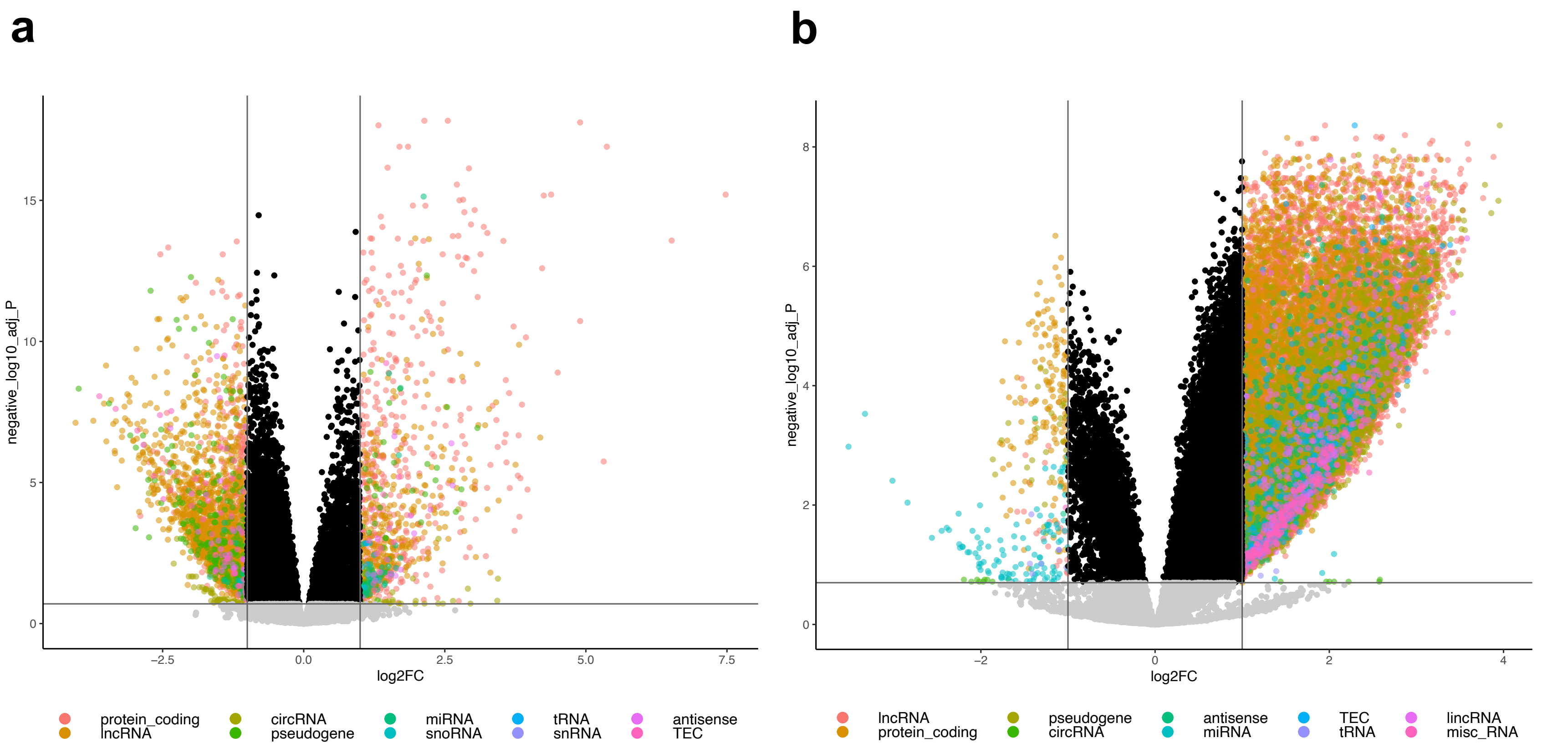


**Supplementary Fig. 1**: Volcano plot showing changes in all transcripts in **(a)** skeletal muscle and **(b)** serum extracellular vesicles (EVs) at h3 vs. pre-exercise in individuals performing TRAD exercise (FDR>0.2, no $log_2FC$ cutoff). Within each biospecimen type these transcripts were pooled with transcripts differentially expressed at other timepoints (h24 for muscle, h0 and h24 for serum EVs) for both exercise doses and summarized into latent variables in the exploratory transcriptomic networks analysis.
