## Supplementary Fig. 2 for "Dynamic Transcriptomic Network Responses to Divergent Acute Exercise Challenges in Young Adults"

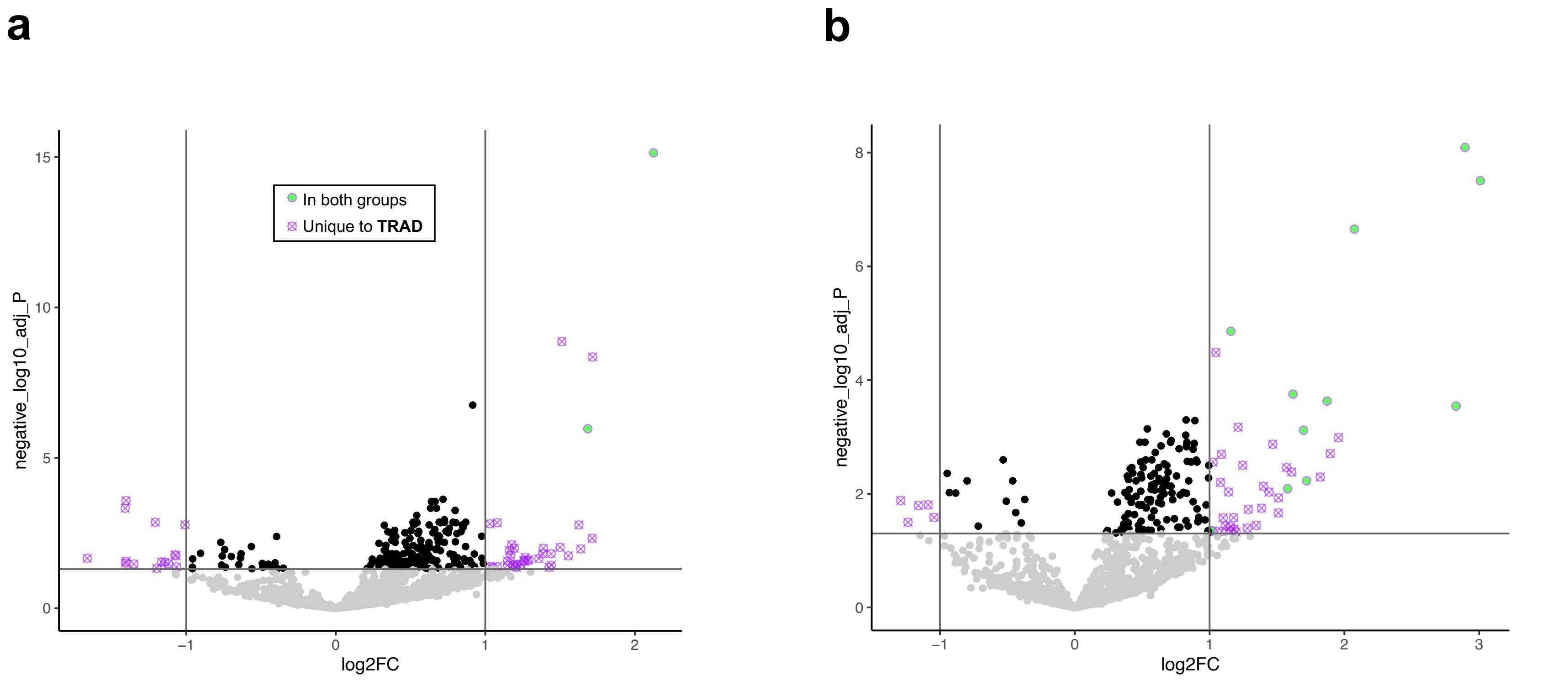


**Supplementary Fig. 2**: Volcano plot showing changes in miRNA expression **(a)** h3 and **(b)** h24 post-exercise in skeletal muscle of individuals performing TRAD exercise (|$log_2$FC| >1, FDR<0.05). Transcripts meeting identical thresholds in HITT are represented as green circles, whereas those shown as crossed-out purple circles were only significant in TRAD (exact values plotted for $log_2$FC and FDR correspond only to TRAD).
