## Supplementary Fig. 3 for "Dynamic Transcriptomic Network Responses to Divergent Acute Exercise Challenges in Young Adults"

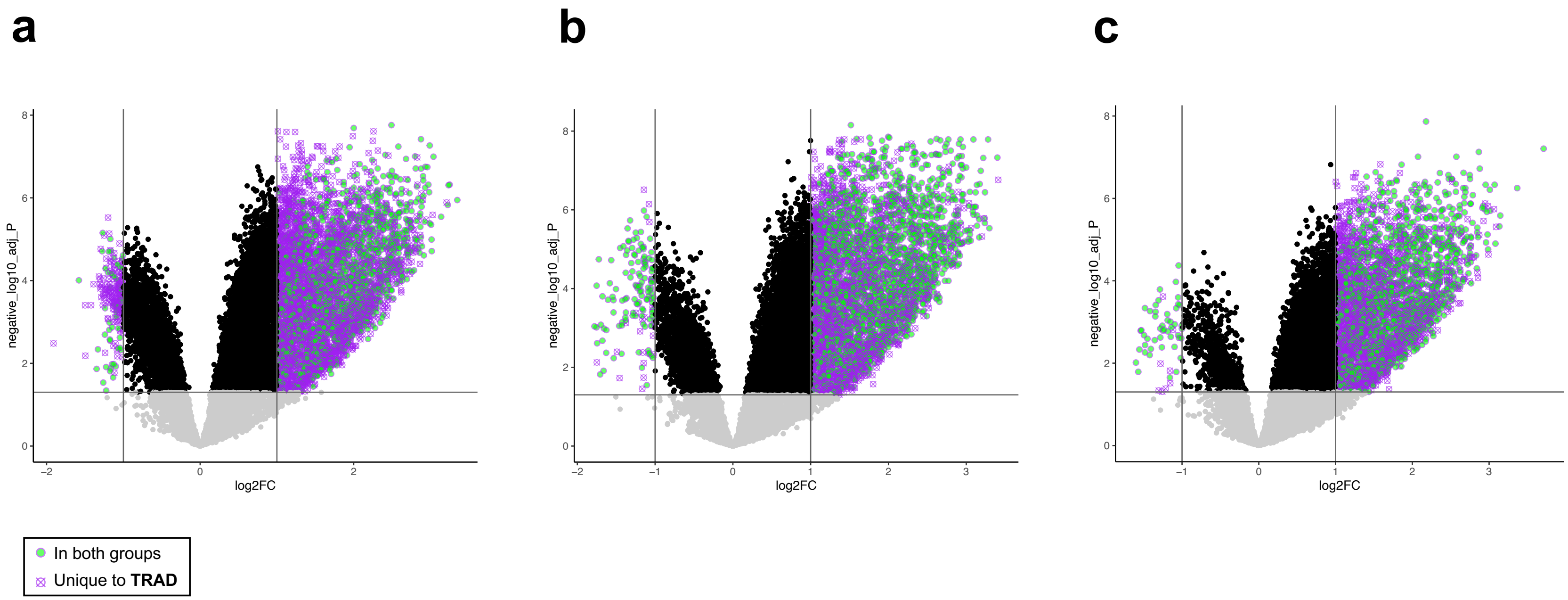


**Supplementary Fig. 3**: Volcano plot showing changes in protein-coding transcript expression **(a)** h0, **(b)** h3, and **(c)** h24 post-exercise in serum extracellular vesicles (EVs) of individuals performing TRAD exercise ($|\log_2 FC| > 1$, FDR<0.05). Transcripts meeting identical thresholds in HITT are represented as green circles, whereas those shown as crossed-out purple circles were only significant in TRAD (exact values plotted for $\log_2 FC$ and FDR correspond only to TRAD).
