## Supplementary Fig. 4 for "Dynamic Transcriptomic Network Responses to Divergent Acute Exercise Challenges in Young Adults"

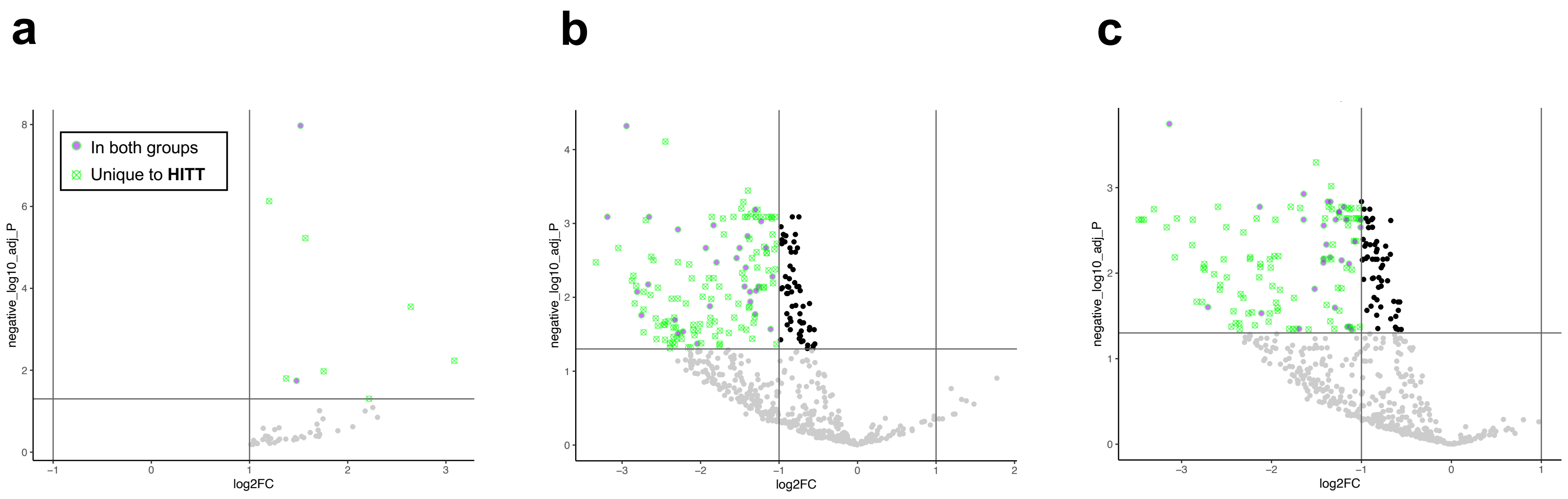


**Supplementary Fig. 4**: Volcano plot showing changes in miRNA expression **(a)** h0, **(b)** h3, and **(c)** h24 post-exercise in serum extracellular vesicles (EVs) of individuals performing HITT exercise. Transcripts meeting identical thresholds in TRAD are represented as purple circles, whereas those shown as crossed-out green circles were only significant in HITT (exact values plotted for $\log_2$FC and FDR correspond only to HITT).
