## Supplementary Fig. 5 for "Dynamic Transcriptomic Network Responses to Divergent Acute Exercise Challenges in Young Adults"

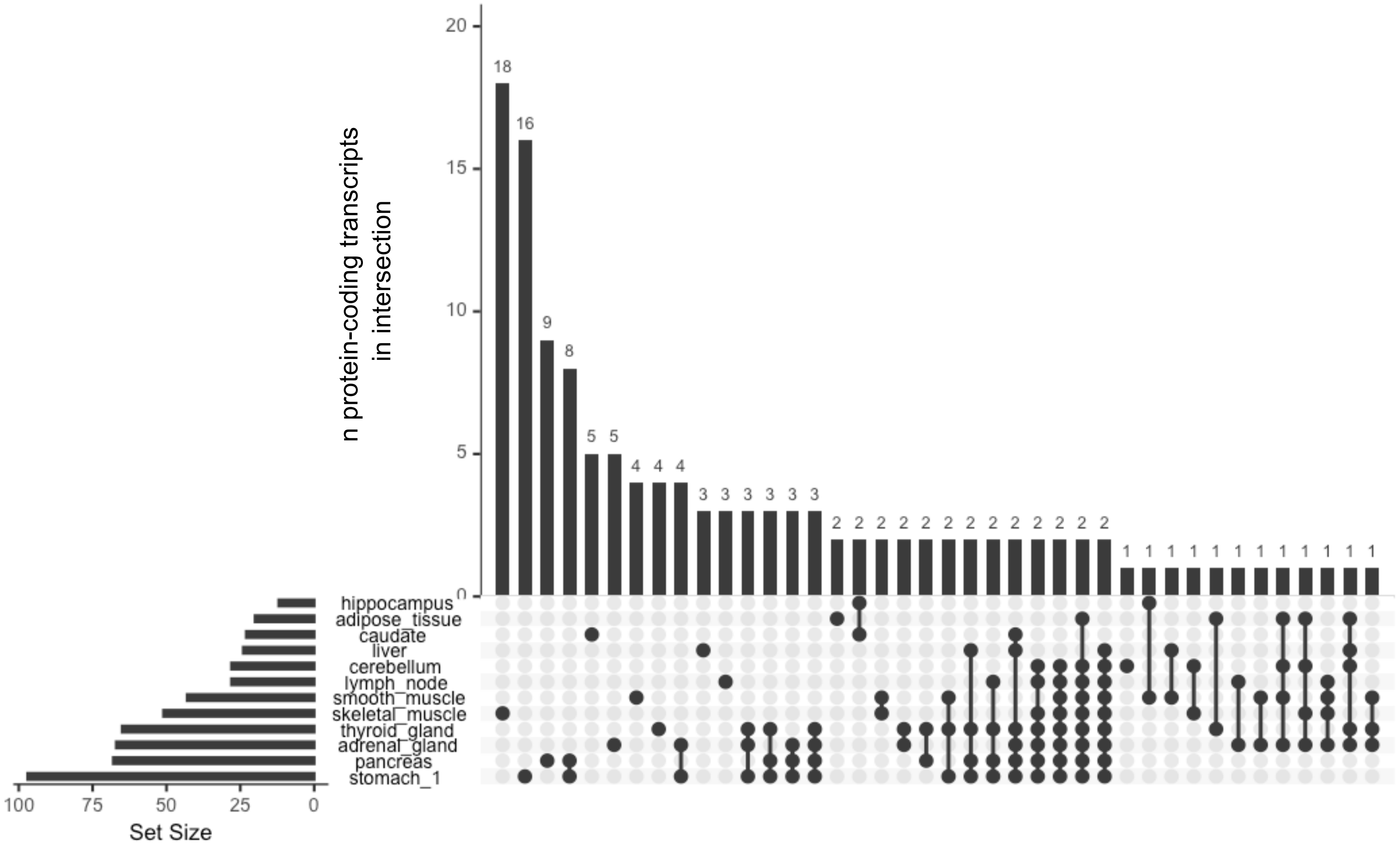


**Supplementary Fig. 5**: UpsetR plot showing tissue enrichment for protein-coding genes in serum extracellular vesicles (EVs) upregulated in both TRAD and HITT at all post-exercise time points, based on the Human Protein Atlas.
